# Androgen receptor contributes to radioresistance through DNA repair and autophagy in AR-positive prostate cancer cells

**DOI:** 10.64898/2025.12.03.690226

**Authors:** Ken-ichi Kudo, Qianhua Feng, Maria Isabel Chosco, Jonathan M. Anzules, Ziyi Huang, Kavya Achanta, Leslie Wenning, Jeshwanth Mohan, June-Wha Rhee, Maedeh Mohebnasab, Evita Sadimin, Zhao V. Wang, Yun Rose Li

## Abstract

Androgen receptor (AR) is a critical therapeutic target in prostate cancer (PCa), and androgen blockade is known to act synergistically with radiation therapy. However, the mechanisms through which AR modulates radiation response are not yet fully understood. In this study, we aimed to investigate the role of AR in mediating radioresistance in PCa. AR-positive LNCaP and castration-resistant C4-2 cells exhibited significantly higher radioresistance than AR-negative cells, as determined by apoptosis and cell viability assays. Following irradiation, most LNCaP cells were arrested in the G1 phase, accompanied by rapid p53 activation and p21 induction. Consistently, AR silencing significantly increased radiosensitivity and reduced DNA-PKcs expression and phosphorylation, suggesting that AR enhances DNA repair, likely through non-homologous end joining (NHEJ). At the cellular level, irradiation markedly induced macroautophagy in LNCaP and C4-2 cells, as evidenced by increased LC3B-II accumulation and autophagic vacuole formation, and the upregulation of 11 autophagy-related genes was identified by whole-transcriptomic analysis. To assess their functional relevance, we performed siRNA-mediated knockdown of selected autophagy-related genes and assessed cell viability and Annexin V/PI staining. Notably, BECN1 and LC3 knockdown significantly enhanced radiosensitivity, with BECN1 knockdown showing an effect comparable to that observed with AR silencing. These results suggest that radiation-induced autophagy promotes the survival of AR-positive prostate cancer cells. Moreover, immunohistochemical analysis of *ex vivo-*irradiated, patient*-*derived PCa tissues from patients with newly diagnosed high*-*Gleason score prostate cancer undergoing prostatectomy further demonstrated that radiation-induced autophagy supports the survival of high-grade AR-positive tumor cells. Collectively, our findings reveal that AR promotes radioresistance in PCa by enhancing both DNA repair and autophagy.

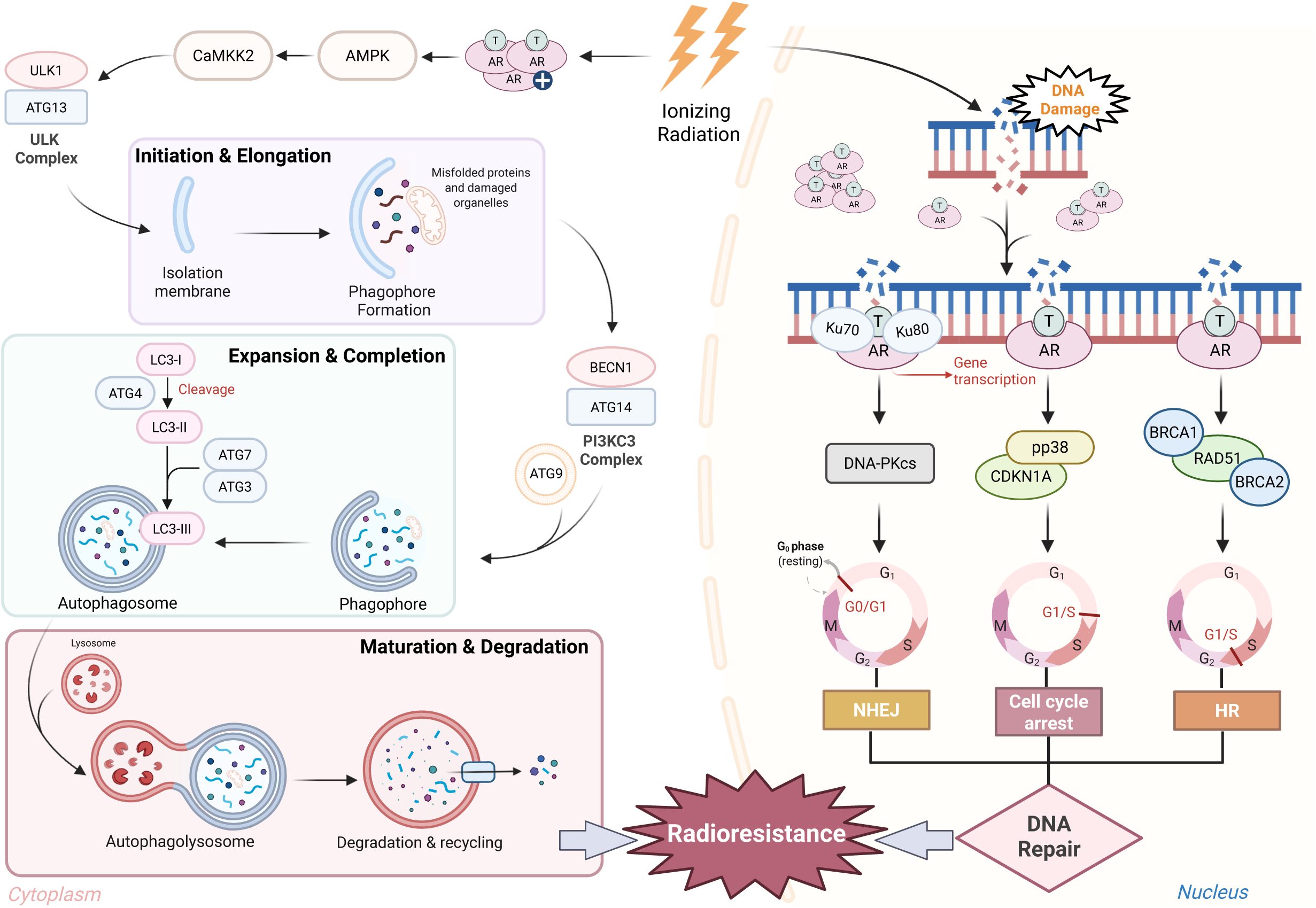

## Introduction

Prostate cancer (PCa) is the second most common cancer among men worldwide and was estimated to cause approximately 35,000 deaths in the U.S. in 2023 (1). The androgen receptor (AR) is a critical therapeutic target for its role in tumor growth and cell survival, primarily through the activation by androgens such as dihydrotestosterone (DHT) (2). Additionally, AR signaling has a further influence on metabolic pathways, including aerobic glycolysis and mitochondrial respiration (3). Androgen deprivation therapies (ADT), which block AR signaling, is widely used as the first-line therapies for PCa for metastatic disease but also as radiosensitizing treatments in combination with radiotherapy for the treatment of aggressive or advanced localized PCa. Unfortunately, over 30% of patients with high risk or node-positive PCa develop recurrences after definitive radiotherapy with ADT, and the vast majority of these patients will ultimately develop resistance to ADT and metastatic disease. Progression of hormone sensitive prostate cancer to castration-resistant prostate cancer (CRPC) is marked by a state in which the tumor cells exhibit overactive AR signaling in an androgen-independent manner (4,5). Several studies have suggested that AR signaling may contribute to DNA damage response (DDR) mechanisms, specifically in repairing radiation-induced double strand breaks (DSBs); therefore, blocking AR signaling can radiosensitize PCa cells (6). However, whether this is the sole mechanism through which AR can modulate radiosensitivity remains unclear, given the broad range of transcriptional targets of AR and the clinical heterogeneity of AR expression in PCa.

In AR-positive models (e.g., LNCaP), AR signaling facilitates repair of radiation-induced DSBs. Prior studies show that activated AR increases DNA-dependent protein kinase catalytic subunit (DNA-PKcs) activity in the non-homologous end joining (NHEJ) pathway, and that pharmacologic AR blockade enhances the cytotoxic effects of radiation (7–9). Consistently, Ku70, which is essential for NHEJ, is reduced with androgen suppression (7). Goodwin et al. reported that activated AR enhances PCa survival post-irradiation in both *in vivo* and *in vitro* experiments by directly modulating DNA-PKcs, an essential mediator of NHEJ repair (8). AR signaling has also been found to promote homologous recombination (HR)-mediated DNA repair (10). Together, these findings suggest that AR activity promotes PCa cell survival after irradiation by enhancing multiple DNA repair pathways.

However, DNA repair may not be the only route by which AR modulates radiation response. Discovered in the 1950s using transmission electron microscopy (TEM), autophagy is the process by which intracellular energy and organelles are renewed through the degradation of intracellular macromolecules, induced by glucose starvation via AMPK activation (11,12). In prostate cancer models, ionizing radiation has been reported to increase autophagy markers, and AR signaling has been linked to the upregulation of autophagy programs that support growth and proliferation (13,14). Functionally, autophagy contributes to radioresistance by limiting the generation of reactive oxygen species (ROS) from dysfunctional mitochondria, clearing radiation-induced damaged macromolecules such as misfolded proteins, and recycling cellular components to sustain the ATP and nucleotide pools required for DNA repair (15,16). Interestingly, AR signaling has also been shown to promote autophagy through its interaction with the AMPK pathway and by regulating autophagic flux (17–19), suggesting that AR may enhance radioresistance not only by promoting DNA repair but also by stimulating cytoprotective autophagy.

In this study, we examine the model in which AR promotes radioresistance through two cooperating mechanisms: enhanced DNA repair and cytoprotective autophagy. We demonstrate that in AR-positive prostate cancer cells, radiation induces autophagy as a prosurvival mechanism, and that blocking autophagy makes cells more sensitive to radiation. Simultaneously, AR activation boosts both DNA repair signaling and autophagic flux. These findings support a therapeutic approach that combines AR pathway inhibition with autophagy blockade to enhance the effectiveness of radiotherapy in AR-positive disease.

## Materials and methods

### Cell lines

In this study, 293T (HEK293T, CRL-3216, RRID: CVCL_0063) and LNCaP clone FGC (LNCaP, CRL-1740, RRID: CVCL_1379) were purchased from ATCC (Manassas, VA). PC3 (RRID: CVCL_0035) and C4-2 (RRID: CVCL_4784) cell lines were provided by Dr. Saul Priceman’s lab and Dr. Sarah Shuck’s lab in Beckman Research Institute of the City of Hope (1500 E. Duarte Rd, Duarte, CA), respectively. LNCaP and C4-2 represent AR-positive prostate cancer models, with C4-2 serving as a castration-resistant derivative of LNCaP, whereas PC3 is AR-negative. HEK293T was included as a non-oncologic, non-prostate epithelial control cell line.

Cells were cultured in RPMI 1640 medium (Thermo Fisher Scientific, Waltham, MA) supplemented with Penicillin/Streptomycin Solution (ATCC), 10% Fetal Bovine Serum (Fisher Scientific, Hampton, NH), HEPES (Thermo Fisher Scientific), and sodium pyruvate (Thermo Fisher Scientific). Unless otherwise specified, RPMI 1640 containing 12.5 mM glucose was used as the medium. In experiments with high and low glucose (HG and LG), all the cells are initially cultured with the 12.5 mM glucose medium first, and then changed to 25 and 5 mM glucose mediums, respectively.

### Irradiation

Cells in dishes were irradiated with X-rays using the X-ray generator MultiRad160 (PRECISION, Madison, CT) at 160 kVp and 25 mA with a 2mmAl filter. The dose rate was 2.8 Gy/min.

### Flow cytometric analysis (FCM)

For autophagy analysis, monodansylcadaverine (MDC) was detected with the Autophagy Assay Kit (Abcam, Cambridge, England), following the manufacturer’s protocol. Afterwards, cells were treated with HG or LG treatment for 48 hours, the medium was exchanged with media containing 20 μM Chloroquine (CQ), and the cells were irradiated with 8Gy via X-rays. 100 nM rapamycin was treated in positive control group. The cells were collected and stained with MDC at 20 hours post-irradiation.

For DDR and apoptotic analyses, FCM measurement with anti-γH2AX and anti-Cleaved Caspase3 (CC3) antibodies (Cell Signaling Technology, Danvers, MA) was performed on irradiated cells. After fixation using 4% paraformaldehyde (PFA) and permeabilization with 0.2% Triton X-100, the cells were treated with the anti-γH2AX and anti-CC3 antibodies for 1 hour on ice. To remove dead cells in the detection of γH2AX signals, Ghost Dye Violet 450 (Tonbo Biosciences, San Diego, CA) was used. These antibodies are listed on Supplementary Table 1.

For cell-cycle analysis, 5-ethynyl-2’-deoxyuridine (EdU) incorporated in genomic DNA was stained using a Click-iT Plus EdU Flow Cytometry Assay Kit (Thermo Fisher Scientific). Cells were treated with the medium containing 10 µM EdU for 30 min before cell recovery, and the EdU was incorporated into S-phase cells during DNA synthesis. After centrifuging at 400 x g for 5 min at 4°C, the cells were washed with D-PBS and subsequently fixed in 4% PFA for 15 min and permeabilized with saponin-based solution. To detect incorporated EdU in DNA strands, the cells were incubated with fluorescent azide for 30 min at room temperature, according to the manufacture’s protocol. Then, cellular RNA was degraded by 100 µg/mL RNase A. Finally, the cells were stained with 10 µM DAPI for 15 min at room temperature. To remove dead cells during FCM, LIVE/DEAD Fixable Dead Cell Stain Kit (Thermo Fisher Scientific) was used.

The apoptotic rate was estimated by Annexin V and propidium iodide (PI) double staining. Cells were incubated in Annexin V binding buffer with Annexin V-AF647 (BioLegend, Inc., San Diego, CA) and PI for 15 min at room temperature in the dark, then analyzed by FCM to distinguish viable and apoptotic cells. The apoptotic rate was calculated based on the percentage of Annexin V/PI double-positive population.

All FCM analyses were performed using LSRFortessa Cell Analyzer (BD Biosciences, Franklin Lakes, NJ, RRID: SCR_018655). Data were analyzed with FlowJo software version 10.8 (BD Biosciences, RRID: SCR_008520).

### Small interfering RNA (siRNA) transfection

siRNA knockdown experiments were performed with siLentFect (Bio-Rad Laboratories). siRNA double strands were synthesized by Thermo Fisher Scientific (Supplementary Table 2). Universal negative control siRNA (NIPPON GENE, Tokyo, Japan) was used as a negative control.

### Cell viability

Cells (1×10^4^) were seeded in a 96-well plate. After 24 hours, cells were irradiated or incubated with the test compounds, including DHT. Each well was measured for absorbance at 492 nm 2 hours after adding the colorimetric CellTiter 96^®^ Aqueous One Solution Cell Proliferation Assay (MTS; Promega, Madison, WI) using a Synergy H4 Hybrid Multi-Mode Microplate Reader (BioTek, Winooski, VT). Cell viability was calculated as a percentage of viable cells vs untreated cells.

### Reverse transcription-quantitative polymerase chain reaction (RT-qPCR)

Total RNA was extracted from cells by using TRIzol reagent (Thermo Fisher Scientific) and was reverse-transcribed into complementary DNA using SuperScript IV VILO Master Mix (Thermo Fisher Scientific), as per the manufacture’s protocol. RT-qPCR was performed using SsoAdvanced Universal SYBR Green Supermix (Bio-Rad Laboratories, Hercules, CA) and QuantStudio Real time PCR system (Thermo Fisher Scientific). *Bax* as a pro-apoptotic marker, *CDKN1A* and *GADD45A* as a cell-cycle arrest marker, *AR* and *PSA* as a prostatic marker, and *ATG2A* as an autophagy marker were measured, respectively. Primer sequences are listed on Supplementary Table 3. The cycling profile included a hot start at 95°C for 120 sec, followed by 35 cycles of 3 steps (95°C of denaturation for 15 sec, 60°C of annealing for 30 sec, 72°C of extension for 30 sec), and a fluorescent signal acquisition at 72°C, with a final dissociation curve analysis. All gene expression values were normalized to β-actin as an internal control.

### TEM

Cells were fixed with 2.5% glutaraldehyde, 100mM Cacodylate buffer (Na(CH_3_)_2_AsO_2_·3H_2_O), pH7.2, at 4°C. Standard sample preparation for TEM was followed including post-fixation with osmium tetroxide, serial dehydration with ethanol, and embedment in Eponate (20). Ultra-thin sections (∼70 nm thick) were acquired by ultramicrotomy, post-stained, and examined on an FEI Tecnai 12 transmission electron microscope (RRID: SCR_022981) equipped with a Gatan OneView CMOS camera (RRID: SCR_014492).

### Immunoblotting

Cells were lysed in complete RIPA buffer containing phosphatase and protease inhibitor cocktails (Nacalai Tesque, Kyoto, Japan). After 30 min on ice, the whole cell lysates were centrifuged at 12,000 x g for 20 min at 4°C and the supernatants were recovered as samples. Protein concentration of each sample was determined by Pierce Rapid Gold BCA protein assay kit (Thermo Fisher Scientific), and equal amounts (20 μg) of protein from each sample was dispensed into 6**–**15% acrylamide gels, separated by SDS-PAGE, and transferred onto Cytiva Amersham Hybond Membranes (Fisher Scientific) or PVDF membranes (Thermo Fisher Scientific). The membrane was incubated with the primary and horseradish peroxidase (HRP)-labeled secondary antibodies. Then, the signal was visualized with Chemi-Lumi One Super (Nacalai Tesque) and detected by ChemiDoc Touch imaging system (Bio-Rad Laboratories, RRID: SCR_019684). All antibodies used in this experiment are listed on Supplementary Table 1.

### Intracellular bead-based multiplex assay (Luminex assay)

To quantitate the phosphorylated proteins relating to DDR accurately, MILLIPLEX_MAP_ DNA Damage/Genotoxicity Magnetic Bead Panel (Millipore Sigma, Burlington, MA) using the Luminex technology was used. LNCaP cells were collected at 1**–**24 hours post-irradiation. Total protein was extracted with complete RIPA buffer, and their concentrations were measured similar to the methods used in western blotting. The samples were filtered using Ultrafree^TM^-CL Centrifugal Filters (fisher scientific) and measured with Luminex FlexMap3D Instrument System (Thermo Fisher Scientific, RRID: SCR_026299), following the manufacture’s protocol.

### RNA sequencing (RNA-seq)

Total RNA was extracted and purified from LNCaP cells at 24 hours post-irradiation by using TRIzol reagent and Direct-zol RNA MiniPrep Plus (Fisher Scientific). RNA quality was evaluated using the RNA ScreenTape assay (Agilent Technology, Santa Clara, CA), resulting in total RNA (> 1.0 μg, OD_260/280_ = 1.8**–**2.2, RIN = 9.1**–**9.8, and 28S/18S = 2.1**–**3.3). For RNA library preparation, Twist RNA Library Preparation kit and Twist Universal Adapter System (Twist Bioscience, South San Francisco, CA) were used. For hybridization, Twist Hybridization and Wash kit with AMP Mix (Twist Bioscience) were used. After hybridization, the quality of cDNA was assessed using the D5000 ScreenTape assay (Agilent Technology). RNA-seq was performed using the NovaSeq6000 (Illumina, San Diego, CA, RRID: SCR_016387).

### Immunocytochemistry (ICC)

Cells were inoculated in glass bottom dishes (Thermo Fisher Scientific) and incubated at 37°C and CO_2_ 5%. After X-irradiation, cells were fixed with 4% PFA for 15 min and permeabilized with 0.2% Tryton-X100 in PBS for 30 min. Nonspecific binding to target sites was blocked using Protein Block (Abcam). Cells were then incubated with primary antibodies at 4°C overnight and with secondary antibodies for 1 hour at room temperature. Finally, cells were treated with DAPI. Images were captured using a confocal microscope LSM900 (Carl Zeiss, Oberkochen, Germany, RRID: SCR_022273). Primary and secondary antibodies used in this study were listed in Supplementary Table 1.

### Neutral comet assay

To quantify the amount of DSBs and evaluate cellular DNA repair, the neutral comet assay was performed using the Trevigen’s CometAssay^®^ Kit (Biotechne R&D Systems) according to the manufacture’s instructions. Cells were collected at 0, 2, and 24 hours after irradiation and diluted to 1.5 × 10^5^ cells/mL, respectively. Diluted cells were combined with LMAgarose at 1:10 (v/v) ratio, spread onto CometSlide™, and lysed overnight at 4°C. Then, the electrophoresis was performed at 21 volts for 45 min at 4°C. Slides were stained with SYBR Gold Nucleic Acid Gel Stain (Thermo Fisher Scientific) for 30 minutes at room temperature. Comets were quantified with AutoComet on GitHub (https://github.com/finkbeiner-lab/AutoComet). The tail length and DNA tail percentage were obtained and used for statistical analyses.

### Patient-derived explant (PDE) culture

Patient-derived PCa tissues were retrieved from the Department of Pathology at City of Hope National Medical Center (City of Hope) to the lab in a timely manner after prostatectomy and divided into 2-3 mm with a sterile knife under sterile conditions. The PCa samples used in this study were derived from patients without previous ADT, and showed Gleason score 4+5=9 tumors. All patient samples were handled according to City of Hope regulations. Tissues were incubated in DMEM/F12 medium (Thermo Fisher Scientific) containing 1 nM DHT and 3 μM Y-27632 at the condition of 37°C and 5% CO_2_. After 1 hour equilibration, explants were irradiated with 0**–**8 Gy X-rays at room temperature and incubated in the same condition for 24**–**48 hours.

### Immunohistochemistry (IHC) staining

PCa tissues on the plate were fixed in 10% neutral-buffered formalin for 48 hours at room temperature and processed for regular paraffin embedding. Immunofluorescence and Hematoxylin-Eosin (H&E) staining for PCa tissues were performed on paraffin-embedded sections. Antigen retrieval was accomplished by autoclaving at 100°C in 10 mM sodium citrate buffer (pH6.0) for 15 min. After blocking with Protein Block, sections were treated with primary antibodies at 4°C overnight. For fluorescent IHC (IHC-F), sections were incubated with fluorescent dye-conjugated secondary antibodies for 1 hour at room temperature and then mounted with VECTASHIELD Antifade Mounting Medium with DAPI (Vector Laboratories, Newark, CA). For bright-field detection, HRP/DAB staining was performed using the Mouse and Rabbit Specific HRP/DAB (ABC) Detection IHC Kit (Abcam), according to the manufacture’s protocol, and sections were mounted with VECTASHIELD PLUS Antifade Mounting Medium (Vector Laboratories). Images were acquired using an LSM900 confocal and a BZ-X810 fluorescence microscopes (Keyence, Osaka, Japan). Quantitative analysis of IHC images was performed using the open-source software QuPath (version 0.6.0-rc4, RRID: SCR_018257). Primary and secondary antibodies used in this study are listed in Supplementary Table 1.

### Statistical Analysis

Statistics were determined using paired Student’s *t* test using Microsoft Excel (Microsoft Corporation, Redmond, WA) and Prism software (GraphPad, SanDiego, CA, RRID: SCR_002798). Data are calculated as the mean ± standard error of the mean (SEM). Statistical significance is represented in figures by: *, P<0.05, **, P<0.01, ***, P<0.001, ****, P<0.0001.

## Results

### AR-positive prostate cancer cell line shows anti-apoptotic properties and superior DNA repair response

To examine whether the underlying mechanisms of radioresistance in PCa cells are dependent on AR and whether protection against DNA damage-related cell death is the primary mechanism of resistance, we first measured the time-dependent radiation responses of LNCaP, PC3, and HEK293T cells using flow cytometry (FCM) (Fig. 1A-C). We measured γH2AX signals to evaluate DNA DSB formation and repair kinetics. Following an 8 Gy dose of irradiation, γH2AX signal levels peaked at 1-hour post-irradiation and declined over the course of 24 hours (Fig. 1D and Fig. S1A–B). However, in LNCaP and PC3 cells, overall γH2AX levels remained higher even at 24 hours post-irradiation compared to those in HEK293T.

**Figure 1.**
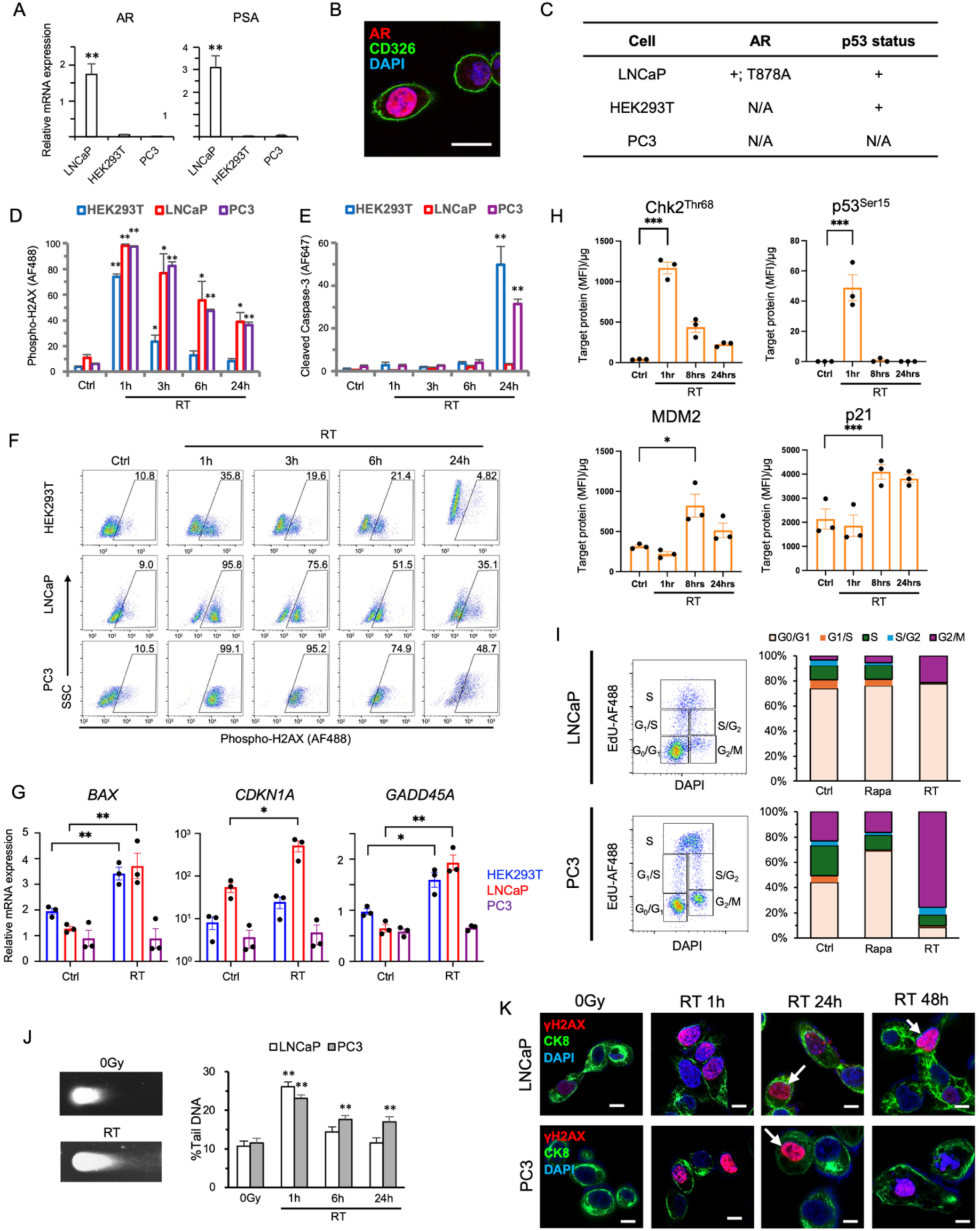
Analysis of DNA damage response and apoptosis induced by post-irradiation. **A,** Relative mRNA expression levels of androgen receptor (AR) and prostate-specific antigen (PSA) in LNCaP, HEK293T, and PC3. All values in the mRNA expression data were scaled to the expression level of β-actin, used as an internal control. **B,** AR expression in LNCaP cells was detected in nucleus by immunocytochemistry (ICC). **C** The gene status is indicated as expression (+) or no expression (N/A) in two types of prostate cancer cell lines and HEK293T. Data are taken from the Cancer Cell Line Encyclopedia (https://depmap.org/portal/ccle/). **D** and **E,** Time-dependent variations of (**D**) γH2AX and (**E**) cleaved caspase-3 (CC3) frequencies generated in HEK293T, LNCaP, and PC3 during 24 hours after X-rays 8Gy radiation-therapy (RT). All flow cytometry (FCM) data represent the mean and SE of three independent assays. **F,** Representative FCM profiles for γH2AX signals induced after irradiation in HEK293T, LNCaP, and PC3 cells. The numbers listed in each panel indicate the γH2AX positive frequencies. **G,** mRNA expression levels of p53-downstream genes in HEK293T, LNCaP, and PC3. **H,** The analysis for radiation response in LNCaP cells exposed with 2Gy by Luminex assay. Superscript letters in each protein name indicate phosphorylation sites. **I,** Analytical results of cell-cycle arrest by EdU assay. Bar plot data represents the frequencies of each population in cell cycle. Rapamycin (Rapa) was used as a positive control for G_1_-phase arrest. **J,** Determination of double-strand breaks (DSBs) generated in LNCaP and PC3 cells after 4Gy exposure by neutral comet assay. The %tail DNA was calculated from the comet tails shown in pictures (n = 100). **K,** ICC data of LNCaP and PC3 stained with anti-γH2AX antibody after 8Gy exposure. White arrow indicates pan-nuclear γH2AX staining. Scale bar = 10μm. *, P<0.05; **, P<0.01; ***, P<0.001 vs. 0Gy or Non-IR by Student’s *t* test.

To further investigate the relationship between AR and radiation-induced apoptosis, we assessed cleaved caspase-3 (CC3) and γH2AX expression. HEK293T and PC3 cells showed statistically significant increases in CC3 24 hours post-irradiation with changes over 30%, while LNCaP CC3 signals remained below 5% (Fig.1E and Fig. S1C). This trend was also reflected in side scatter (SSC) measurements, which are associated with nuclear fragmentation during apoptosis (Fig. 1F) (21). HEK293T and PC3 cells showed increased SSC post-irradiation; SSC levels remained low in LNCaP cells (Fig. 1F, Fig. S1D). Consistent with their reduced apoptotic response, LNCaP cells displayed a distinct new γH2AX-positive population at 24 hours post-irradiation (Fig. 1F and Fig. S1B), indicating a sustained DNA damage response rather than apoptotic fragmentation. RT-qPCR results indicated that LNCaP cells exhibited the highest expression levels of p53-downstream genes, such as BAX, following irradiation (Fig. 1G). The Luminex assay data demonstrated early activation of the p53 pathway within a few hours post-irradiation (Fig. 1H). Together, these results indicate that LNCaP cells are resistant to radiation-induced apoptosis.

We next performed cell-cycle analysis using EdU to compare radiation-induced cell-cycle arrest in two prostate cancer cell lines (Fig. 1I). Before RT, both LNCaP and PC3 cells were predominantly in the G_0_/G_1_ phase. After 20 hours post-irradiation, cells entered the G2/M checkpoint, especially in PC3 cells. Notably, the predominance of G_0_/G_1_ phase population persisted in LNCaP cells, suggesting that LNCaP cells are preferentially engaged in the NHEJ pathway to repair radiation-induced DSBs. Consistent with these findings, RT-qPCR and Luminex assay results showed that LNCaP cells demonstrated increased expression of *p21^WAF1/CIP1^*(*CDKN1A; Fig. 1H*), which is involved in G_1_-S arrest (23), and *GADD45A (Fig. 1E)*, which is associated with G_2_ arrest post-irradiation (24). In contrast to the LNCaP results, PC3 cells showed no reaction (Fig. 1G), consistent with the conventional findings that PC3 cells have a p53 loss of function. Neutral comet assay results demonstrated that radiation-induced DSBs were more rapidly repaired in LNCaP cells than in PC3 cells (Fig. 1J). In addition, confocal fluorescent images showed pan-nuclear γH2AX signal, which correlates with DNA-PKcs activity (22), in LNCaP cells up to 48 hours post-irradiation (Fig. 1J). These findings were consistent with our results showing that γH2AX signals remained high in LNCaP cells even 24 hours after irradiation and that a distinct new population emerged, as shown in the γH2AX results (Fig. 1D and S1B). Taken together, these observations suggest that LNCaP cells have a rapid radiation response and superior DNA repair ability.

### Androgen receptor (AR) influences cell proliferation and the expression of DNA repair proteins

To analyze the underlying molecular mechanisms linking AR to radioresistance, we suppressed AR activity using siRNA knockdown and the signaling inhibitors enzalutamide and abiraterone acetate (23,24). On average, we found that siRNA knockdown significantly reduced *AR* expression levels by 75% when compared to the scramble siRNA control group (Fig. 2A). To observe the effects of AR knockdown and signaling inhibition, we measured *KLK3/PSA* expression levels with RT-qPCR, and found that *KLK3/PSA* expression levels also decreased. In enzalutamide-treated and abiraterone acetate-treated groups, *AR* expression remained unchanged, but *KLK3/PSA* expression levels decreased significantly. No significant difference was observed in the expression of p53-downstream genes across all groups (Fig. 2A and Fig. S2A). Interestingly, *AR* expression was also significantly increased 20 hours after irradiation.

**Figure 2.**
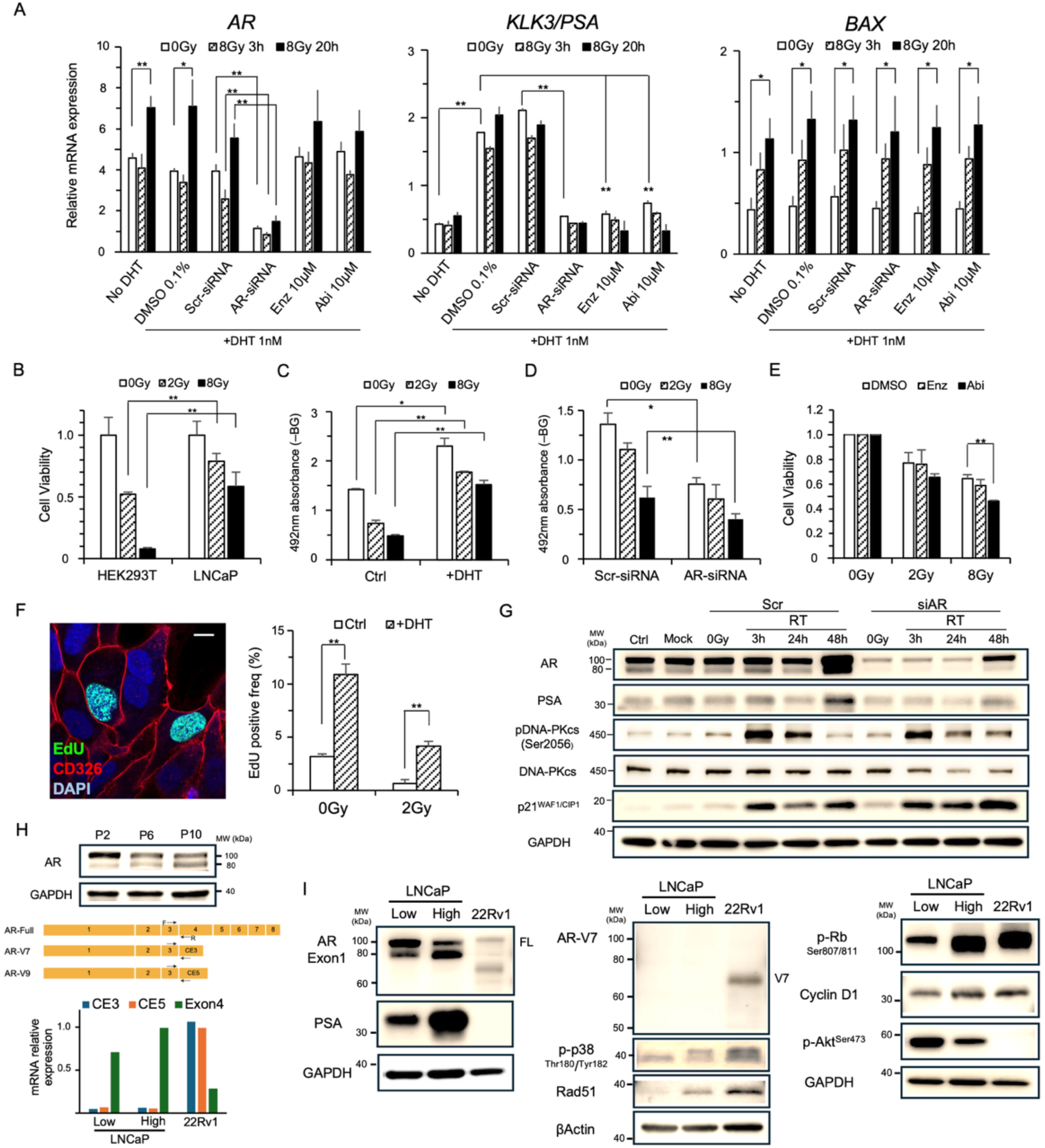
The results of AR siRNA knockdown and AR signaling inhibition. Experiments were conducted with medium containing charcoal-filtered FBS. **A,** Relative mRNA expression levels of *AR*, *KLK3/PSA*, and *BAX* genes in LNCaP cells. Enz and Abi refer to enzalutamide and abiraterone acetate, respectively. The data demonstrates the fold difference of each mRNA expression against negative controls and was obtained from the means and SEs of three independent assays. **B–E,** Cell viabilities and 492nm absorbance data indicate the comparison between (**B**) HEK293T and LNCaP cells, and for LNCaP only: (**C**) With and without dihydrotestosterone (DHT), (**D**) Scramble (Scr)-siRNA and AR-siRNA, (**E**) Enz or Abi-treated groups after irradiation. **F,** Average EdU frequencies from the ICC data stained with fluorescent azide targeting EdU and anti-CD326 antibody, and total counts reach more than 100 per column. Scale bar = 10μm. **G,** Western blotting analysis for Scr and AR-knockdown groups. LNCaP cells past passage number 9 were treated with 1 nM DHT for 48 hours and then irradiated with 8Gy of X-rays. The numbers on the vertical axis represent molecular weight (kDa). **H,** Comparison of AR protein expressions and mRNA expressions for cryptic exons (*CE3* and *CE5*) in the AR gene between passage numbers. The low and high mRNA expression graphs indicate low (P5) and high (P9) passage numbers. **I,** Comparison of each protein expression between low (P4) and high (P10) passage numbers in LNCaP cells. *, P<0.05, **, P<0.01 by Student’s *t* test.

To examine the effect of AR on cell proliferation, we measured cell viability and absorbance at 492nm using the MTS assay, a colorimetric assay that quantifies metabolically active cells by converting MTS to a soluble formazan product. We found that cell viability post-irradiation was higher in LNCaP than in HEK293T cells (Fig. 2B), which is consistent with the CC3 results (Fig. 1E). Androgen-supplemented groups demonstrated higher cell proliferation than that in control groups, even after irradiation (Fig. 2C,F, and Fig. S2B–C). In contrast, this proliferative activity was lost after AR was knocked down (Fig. 2D). Similarly, we observed a significant reduction in cell viability following AR signaling inhibition with abiraterone acetate. (Fig. 2E). Taken together, these results suggest that in the AR-expressing groups, the complete arrest of the cell cycle does not occur, proliferative activity continues even after irradiation, and the transcriptional activity of AR affects radioresistance in LNCaP cells.

To explore how AR might influence DNA repair pathways, we examined the relationship between AR and DNA-PKcs, which has been previously implicated in AR-driven radioresistance. Western blot analysis showed that the expression and phosphorylation of DNA-PKcs were inhibited after AR knockdown (Fig. 2G). In the high-passage LNCaP cells, a secondary AR band appeared near 80 kD (Fig. 2H). Interestingly, the abundance of this band increased with passage number, and it tested negative for known AR variants, including AR-V7 and -V9 (Fig. 2H-I and Fig. S2D). Notably, PSA and Cyclin D1, which are well-established AR transcriptional targets, were elevated in high-passage LNCaP cells (25) (Fig. 2I). Because these genes directly report AR transcriptional output, their upregulation indicates that AR activity increases over time and supports the idea that high-passage LNCaP cells develop more ligand-independent and clinically relevant AR-driven behavior. Phosphorylation of Rb, a marker for G_1_-S phase transition that is downstream of Cyclin D1, was similarly elevated in high-passage LNCaP cells (26–28). Lastly, we found that AR at high passages also promoted the expression of DNA repair genes, such as Rad51, and increased the phosphorylation of p38, a key component of HR repair (Fig. 2I). These findings support the hypothesis that AR activity drives cell proliferation and DNA repair. In contrast, Akt phosphorylation levels decreased at high passage numbers, contrary to what is typically expected for AR-driven signaling.

### Radiation induces autophagy in AR-positive prostate cancer cell line

From our previous analysis, we identified a reduction in *ATG2A* expression in AR knockdown and abiraterone-treated groups (Fig. S2E). To determine whether autophagy occurred after irradiation in AR-expressing cells, we treated cells with chloroquine (CQ). CQ inhibits lysosomal and autophagosomal fusion, thereby delaying autophagy and promoting the accumulation of autophagic vacuoles (AVs) and markers such as LC3B-II within the cell (Fig. 3A) (29). We used MDC, a traditional, non-specific dye that accumulates in acidic vesicles, to assess AV formation (30). We found that MDC signals in LNCaP cells increased significantly post-irradiation, even exceeding the levels observed under rapamycin treatment (Fig. 3B and Fig. S3A). While HEK293T and PC3 did not see any significant increase in MDC, we did see an increase in *ATG2A* mRNA expression in HEK293T cells (Fig. S3B). Given that autophagy is governed by AuTophaGy-related genes (ATGs) (31), we comprehensively investigated the mRNA expression levels of ATGs in LNCaP cells with RNA-seq analysis. Consistent with previous results, ATG mRNA demonstrated significant increases in expression, including in *ATG2A/B*, *ATG13*, *ULK1*, *BECN1*, *ATG7*, *ATG9A*, *ATG4A/B*, and *ATG14* post-irradiation (Fig. 3C).

**Figure 3.**
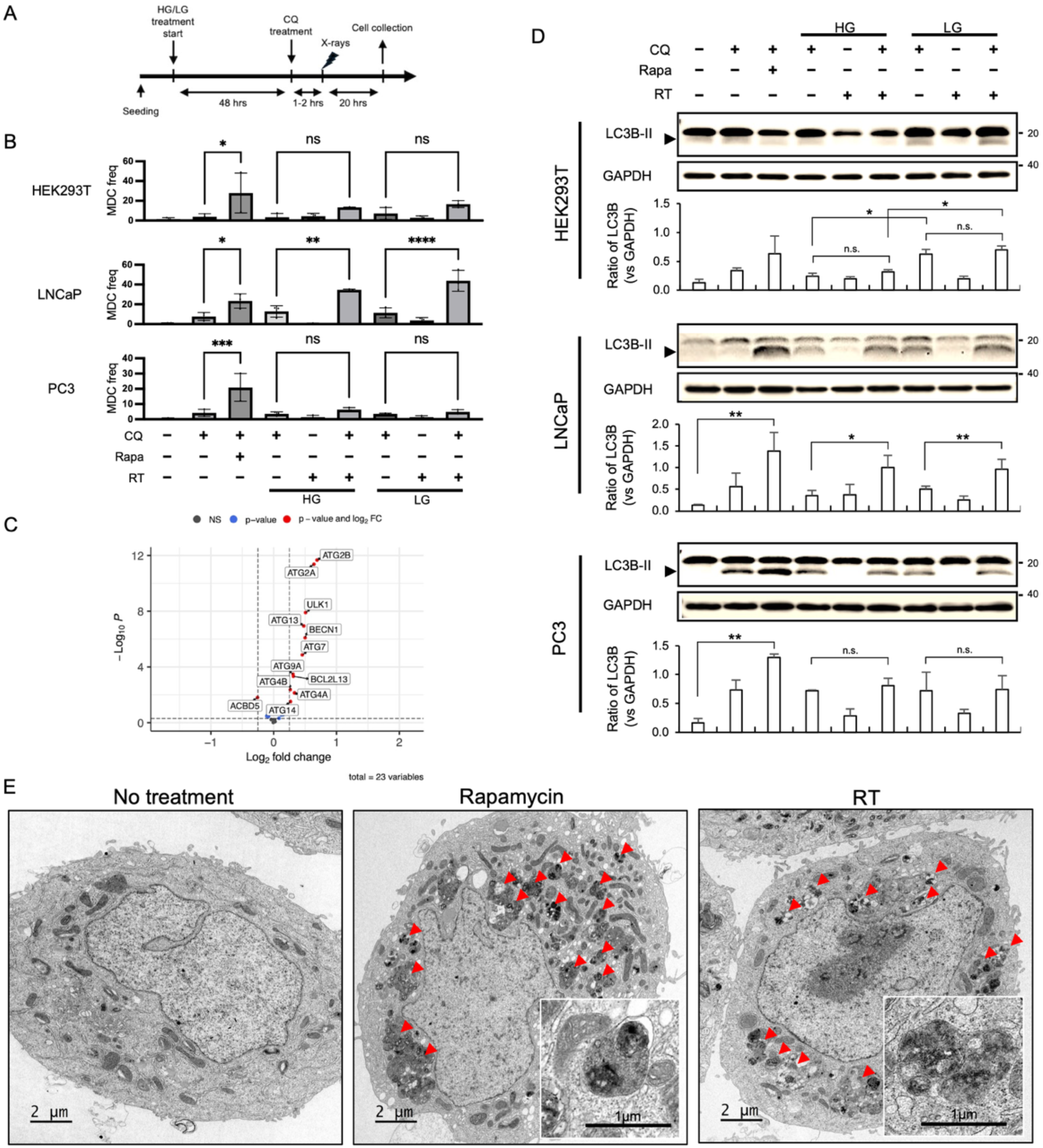
X-irradiation can strongly induce autophagy in AR-positive cells. **A,** Experimental procedure for analyzing autophagy caused by irradiation. **B,** Monodansylcadaverine (MDC) frequencies were estimated in each cell line with FCM. All data represent the means and SEM of three independent assays. CQ refers to chloroquine, and Rapa refers to rapamycin. *, P<0.05, ***, P<0.001 vs. negative control by Student’s *t* test. **C,** Volcano plot of RNA-seq data showing up/downregulation of differentially expressed autophagy-related genes when comparing unirradiated (0 Gy) to irradiated (8Gy) conditions. **D,** LC3B-II ratios were measured with western blot analysis to detect autophagy. GAPDH was used to estimate ratios as an internal control. *, P<0.05, **, P<0.01 by Student’s *t* test. **E,** Transmission electron microscopy images of LNCaP cells treated with rapamycin and 8Gy irradiation, obtained from the MDC assay. Red arrows indicate autophagic vacuoles. Scale bar = 2μm.

Next, we quantified LC3B (ATG8) protein levels, which play a critical role in autophagosome biogenesis and maturation (32). Consistent with MDC results, we did not observe any differences in the LC3B-II/GAPDH ratio between the irradiated and unirradiated groups in HEK293T and PC3 cells. However, in LNCaP cells, we measured significant increases at 20 hours post-irradiation, comparable to the rapamycin-treated group (Fig. 3D). In addition, consistent with previous results, we directly observed AV formation within LNCaP cells using TEM (Fig. 3E) and found AV levels to be comparable to those in rapamycin-treated group in LNCaP cells 20 hours post-irradiation, which were also determined to be late/degradative AVs (33). In HEK293T cells, LC3B-II/GAPDH ratios and phosphorylated AMPKα (p-AMPKα) levels were higher in the LG-treated group than that in the HG-treated group (Fig. 3D and Fig. S3D-E). These findings suggest that LG treatment may induce autophagy signaling via the p-AMPKα pathway, consistent with previous reports (11,12,34). Phosphorylated p38 (p-p38), which is known to support cell survival and DNA repair (35,36) and influence the MAPK pathway, was also elevated in LG-treated cells after irradiation (Fig. S3D). These findings explain the γH2AX results of HEK293T shown in Fig.1D that LG condition may be favorable for DNA repair and survival due to p-p38 augmentation in normal cells.

### BECN1 and MAP1LC3B may be novel therapeutic targets in radiation therapy against AR-positive prostate cancer including CRPC

Previously, we found that ATG expression increased after irradiation and that autophagic flux was elevated, as indicated by LC3-II ratios and AV analyses. Therefore, we next selected three main genes in the autophagic process, BECN1 and ATG2A, which represent distinct processes of autophagy, and MAP1LC3B, from the 11 genes identified in the RNA-seq analysis (Fig. 3C and 4A). We then performed siRNA-mediated knockdown on C4-2 cells to evaluate the potential of ATG genes as therapeutic targets in radiation therapy against PCa in the LNCaP-derived CRPC cell line model (Fig. 4).

**Figure 4.**
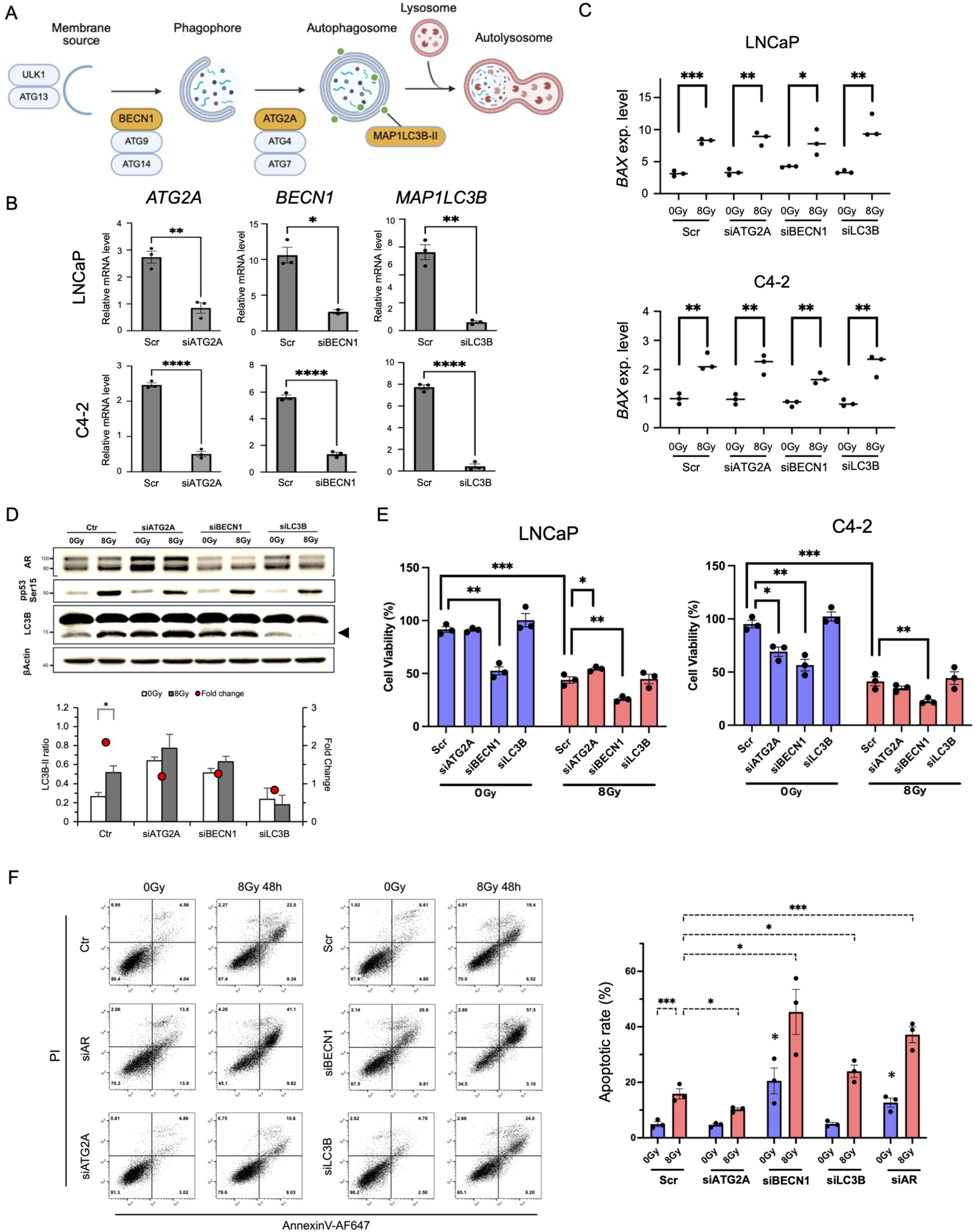
Effects of ATG-siRNA knockdown on radiosensitivity in AR-positive prostate cancer cell lines. Experiments were performed using medium supplemented with charcoal-stripped FBS. **A,** Schematic diagram showing the autophagic process, including nine ATG genes that were upregulated in response to radiation, as identified in Fig.3. **B-C,** Relative mRNA expression levels of *ATG2A*, *BECN1*, and *MAP1LC3B* **(B)**, and *BAX* **(C)**, a proapoptotic and p53-downstream gene, in LNCaP and C4-2 cells. mRNA expression was normalized to *GAPDH*. Data represent the means and SE from three independent experiments. **D,** Western blotting analysis of ATG-siRNA knockdown groups. Molecular weights (kDa) are indicated on the left. The arrow on the right indicates LC3-II. The graph shows LC3B-II/β- Actin ratios and the fold change between 0Gy and 8Gy. **E,** Cell viability of LNCaP and C4-2 after ATG gene knockdown. **F,** Apoptotic rates, represented by Annexin V/PI double-positive frequencies, were measured in C4-2 cells using flow cytometry (FCM) with Annexin V-AF647 antibody and PI. Data represents the means and SE from three independent experiments. *, P<0.05; **, P<0.01; ***, P<0.001. Significance marks without bars in the 0Gy condition of siBECN1 and siAR groups indicate comparisons against 0Gy in the Scr group.

Following siRNA knockdown, mRNA expression of the target genes was significantly reduced to approximately one-third to one-eighteenth of the control levels in LNCaP and C4-2 cells (Fig. 4B). Next, we measured the expression of *BAX*, but we did not find any significant differences between the control and knockdown groups (Fig. 4C). Although *BAX* expression was slightly lower in the irradiated siBECN1 group compared to others, the difference did not reach statistical significance (p = 0.06).

To assess the effects of ATG knockdown on autophagy, we analyzed LC3B-II protein levels in C4-2 cells (Fig. 4D). In the control group, the LC3B-II/β-Actin ratio significantly increased after irradiation, whereas the LC3B-II/β-Actin ratio was unchanged in the ATG-knockdown groups. The results confirm that radiation-induced autophagy was inhibited by ATG knockdown. In addition, the siATG2A and siBECN1 groups exhibited relatively high basal LC3B-II ratios, and we detected AR and phospho-p53 (Ser15, pp53). The pp53 levels demonstrated no significant differences between groups, consistent with *BAX* mRNA expression (Fig. 4C). AR expression in C4-2 cells appeared as a doublet, consisting of the intact ∼100kDa band accompanied by an ∼80kDa band, like that observed in LNCaP cells.

After confirming that ATG knockdown suppressed autophagy, we evaluated the radiosensitivity of LNCaP and C4-2 cells using cell viability assays. In both cell lines, siBECN1 treatment significantly reduced cell viability 48 hours after knockdown (Fig. 4E). In addition, knockdown of ATG2A in C4-2 cells resulted in a significant decrease in cell viability in the unirradiated group. In contrast, the irradiated LNCaP cells in the siATG2A group exhibited a significant increase in cell viability compared with the control group. Since cell viability reflects the total cell number, these results likely include cells that underwent apoptosis or cell-cycle arrest because of DDR after irradiation. To further evaluate radiation-induced apoptosis in C4-2 cells, we performed FCM to detect Annexin V/PI double-positive cells. We found that apoptotic rates were markedly increased in the siBECN1 group at 48 hours after knockdown, even in the absence of irradiation, consistent with our results shown in Fig. 4E (Fig. 4F). The extent of this increase was comparable to the rise observed in the siAR group (siBECN1_8Gy_ vs siAR_8Gy_, n.s.).

Meanwhile, the siLC3B group also showed a significant increase in apoptotic cells after irradiation compared with the control group, although the magnitude of this increase was smaller than that observed in the siBECN1 group. In contrast, the siATG2A group demonstrated a significant decrease in apoptosis (p < 0.05). Taken together, these findings suggest that inhibiting autophagy via BECN1 and LC3B enhances radiosensitivity in AR-positive prostate cancer cells.

### *Ex vivo* validation of AR-associated autophagy and radioresistance in patient-derived prostate cancer tissues

To evaluate the clinical relevance of these *in vitro* findings, we performed IHC analysis using a patient-derived explant (PDE) model to examine AR expression, radiation-induced autophagy markers, and their association with radioresistance. Tissues were collected at the time of clinically indicated radical prostatectomy from patients with newly diagnosed high-risk prostate cancer (Gleason 8-10). Among the four human PCa tissues used in this study, three exhibited positive expression of AR, PSA, and CK8, consistent with a prostate luminal epithelial phenotype (Fig. 5A–B and Fig. S4A–D). As expected, we observed the nuclear translocation of AR in both benign and malignant prostate tissues following treatment with 1 nM DHT (insets in Fig. 5B), and the protein expression levels of AR and PSA remained stable up to 72 hours after irradiation (Fig. S4A–D). PCa tissues displayed distinct γH2AX foci at one hour post-irradiation, and we also observed pan-nuclear γH2AX staining (Fig. S4E). These findings indicate that the patient-derived prostate cancer tissues used in this study are maintained adequately during PDE culture for 72 hours, preserving AR activity and exhibiting radiation responses, comparable to those observed in *in vitro* experiments.

**Figure 5.**
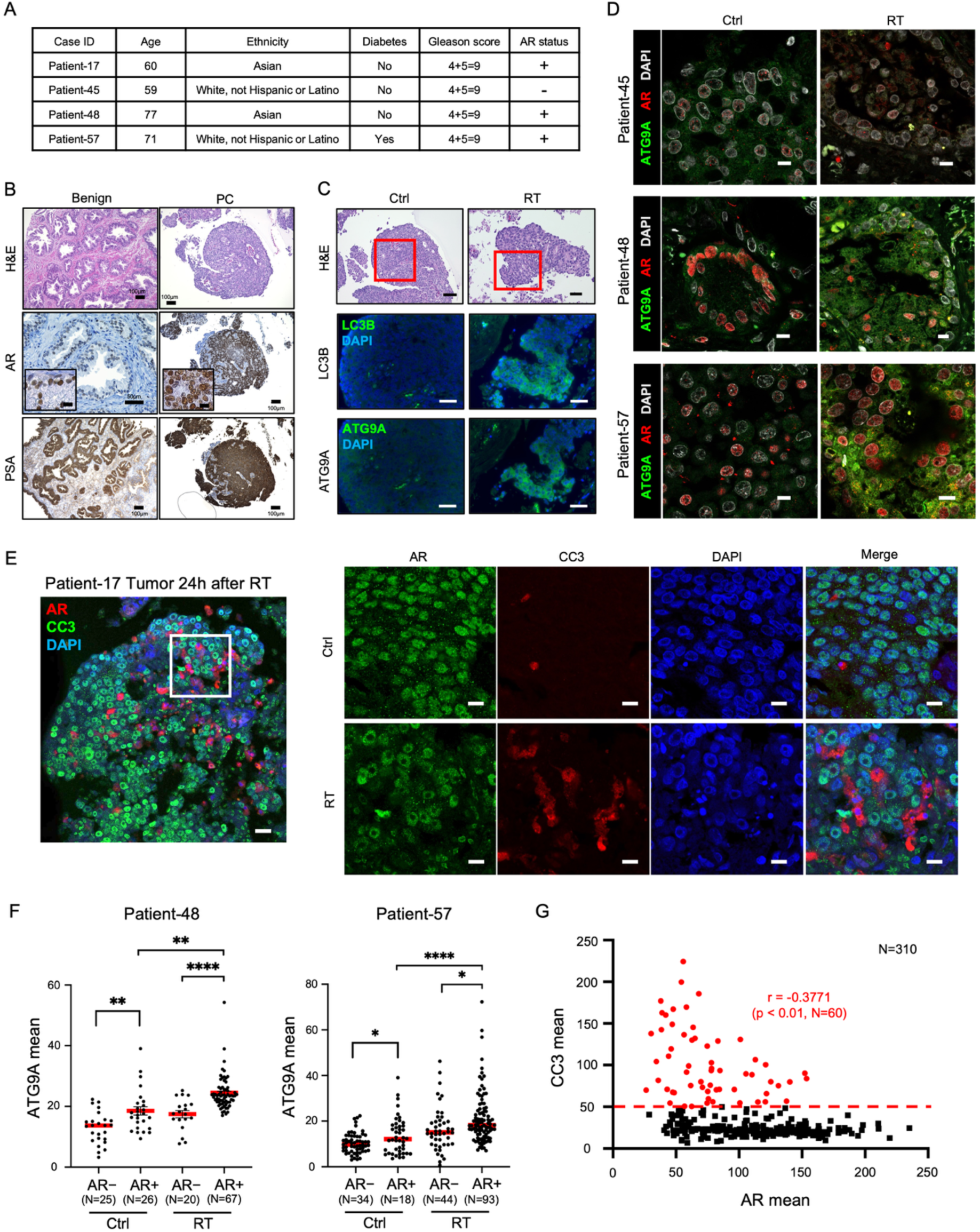
Immunohistochemistry (IHC) analyses to measure radiation responses in patient-derived explants. **A,** Patient samples were obtained from individuals designated as Patient-17, −45, −48, and −57, and the clinical information acquired included age, race, diabetes status, Gleason score, and AR expression status. **B,** Histology of benign and prostate cancer (PCa) tissue in Patient-17 as a representative sample. Tissues were cultured with DMEM/F12 medium containing 1 nM DHT and 3μM Y-27632 within 1 hour after prostatectomy. Representative images include hematoxylin and eosin (H&E) and IHC for AR and PSA. Scale = 10μm in the inset of AR. **C,** LC3B and ATG9A fluorescent images of PCa tissues from Patient-17. Red boxes in H&E images indicate the field of view for fluorescent images. Scale bar = 50μm. **D,** Confocal fluorescent images of AR and ATG9A in Patient-45, −48, and −57. Because the LC3B-II methodologies are not applicable to formalin-fixed patient tissues, ATG9A was analyzed as an alternative autophagy-related marker. Scale bar = 10μm. **E,** Confocal fluorescent images for AR and CC3 in Patient-17. White box in left image represents the field of view for fluorescent images. Scale bar = 10μm. **F,** Quantification of ATG9A intensity in AR-positive and AR-negative cells from Fig.5D (Patient-48 and −57). AR signal threshold was set at a relative intensity using QuPath software, and cells with above or below this threshold were defined as AR+ and AR−, respectively. Red bars indicate median intensity in each group. **G,** Scatter plot showing an inverse correlation between AR and CC3 obtained from images in Fig.5E. Each dot represents a cell; the x- and y-axes correspond to mean fluorescence intensities of AR and CC3, respectively, quantified using QuPath. Red line indicates the threshold for CC3 positivity. Ctrl: 0Gy, RT: 8Gy.

At 24–48 hours post-irradiation, strong expression of autophagy markers LC3B and ATG9A was observed in AR-positive tumor regions, which were not detected in AR-negative samples (Fig. 5C–D). Quantitative analysis further revealed that ATG9A expression levels were significantly higher in AR-positive cells than in AR-negative ones (Fig. 5F and Fig. S4F). The observed rise of autophagy markers post-irradiation, coupled with our previous results (Fig. 3C–D), demonstrates that autophagy is induced in AR-positive prostate cancer cells *in vivo* following irradiation. Finally, to investigate the relationship between AR expression and radiation-induced apoptosis, we analyzed AR/CC3 double-stained sections to identify apoptotic cells within prostate cancer tissues after irradiation. We found that AR-positive cells exhibited low CC3 signals, and AR and CC3 expression levels were significantly inversely correlated (Fig. 5E and G; p-value < 0.01). Collectively, these results suggest that high-grade AR-positive prostate cancer cells may evade radiation-induced apoptosis by inducing autophagy.

## Discussion

Despite many decades of clinical data supporting the benefit of combining AR inhibition with radiotherapy to improve curative outcomes of men with aggressive, localized prostate cancer, the fundamental mechanisms through which AR inhibition results in radiosensitization remain unclear. In this study, we examine the mechanisms through which AR modulates radioresistance in PCa cells. Contrary to the expectation that hormone-sensitive or AR-expressing PCa cells would be more radiosensitive, LNCaP cells exhibited a strong resistance to radiation-induced apoptosis when compared to HEK293T and PC3 cells. Following irradiation, LNCaP cells were arrested in the G_1_ phase, as indicated by elevated p21 expression, a downstream target of p53. This is consistent with efficient DSBs repair via the NHEJ pathway and thus increased repair-mediated radioresistance. As autophagy has previously been implicated in both radioresistance and radiosensitization, we sought to resolve this question by evaluating how AR may participate in RT-induced autophagy. Indeed, radiation-induced autophagy was observed only in the two AR-positive cell lines, LNCaP and C4-2, whereas post-irradiation autophagy could not be observed in AR-negative cell lines.

To test whether AR expression directly contributes to radioresistance, we knocked down AR and found that AR knockdown was associated with reduced cell proliferation and reduced expression of DNA repair proteins, such as DNA-PKcs and Rad51, supporting our observation that LNCaP cells exhibit a high DNA repair capacity. In addition, these changes were prominent in cells expressing an AR C-terminus-truncated form, which is believed to be associated with increased AR transcriptional activity. Classically, after androgens such as DHT bind to AR-LBD, AR translocates into the nucleus and binds to androgen response elements (AREs), thereby enhancing target gene expression (2). However, some studies have suggested an alternative non-canonical pathway in which partial ubiquitination and degradation of the C-terminal—possibly via Ser-515 phosphorylation—generate a truncated AR centered on the N-terminal domain (NTD) (37–39). This truncated AR isoform can enter the nucleus more rapidly and activate AREs independently of ligand binding. Our findings suggest that truncated AR may be a driver of radioresistance in early CRPC, where AR is still expressed and cells remain hypersensitive to testosterone, rather than truly androgen-independent.

Additionally, we observed an increase in AR mRNA expression 20 hours post-irradiation, followed by a subsequent increase in AR protein levels by 48 hours. Several studies have reported that AR expression is upregulated in response to radiation or ROS exposure (40,41). Notably, p21 levels remain elevated at 48 hours post-irradiation. This sustained expression raises several questions, as it may indicate prolonged G_1_ phase arrest or persistent cellular stress signaling. Both of these can contribute to increased radioresistance. Given the established role of AR as a therapeutic target for radiosensitizing PCa, it is essential to investigate further the mechanisms underlying radiation-induced AR upregulation and their implications for treatment strategies.

Because radiation induces autophagy, we investigated whether ATG genes could modulate radiosensitivity in AR-expressing PCa cells. BECN1 plays a central role in initiating the autophagy process (42). In our study, siRNA-mediated BECN1 knockdown reduced PCa cell viability and enhanced radiosensitivity in C4-2 cells, an AR-positive, castration-resistant prostate cancer cell line commonly used as a CRPC model. These findings indicate that autophagy contributes to cell survival and radioresistance in prostate cancer. Moreover, Beclin-1–dependent autophagy has been implicated in therapy response in CRPC, with modulation of Beclin-1 and the AR–Beclin-1 complex altering sensitivity to enzalutamide, and repression or inhibition of Beclin-1–mediated autophagy resensitizing prostate cancer cells to docetaxel and other agents (43,44).

Moreover, others have reported increased drug sensitivity to enzalutamide and other agents following BECN1 suppression, and our results align with these findings, further emphasizing the potential therapeutic role of BECN1 as a target in PCa, including CRPC.

Additionally, our findings suggest that MAP1LC3B may be a potential therapeutic target. Other ATG genes, such as ULK1/2, which have already been reported as AR-regulated targets (14), may also enhance radiosensitivity in prostate cancer when their expression is downregulated, likely by inhibiting autophagy. Given that p53-downstream gene transcription was highest in the RT-PCR results, AR signaling in LNCaP and C4-2 cells may synergize with p53-mediated suppression of mTOR (45,46). Consistent with these *in vitro* findings, irradiated human PCa tissues showed increased ATG9A expression, a key regulator of autophagy, and an inverse correlation between AR signaling and CC3. These observations suggest that AR contributes to radioresistance by promoting DNA repair and autophagy.

Although siRNA-mediated MAP1LC3B knockdown did not significantly reduce cell viability compared with the scrambled control, this may reflect the concomitant suppression of mitophagy, in addition to bulk autophagy, when LC3B-II is reduced (47). Under these conditions, mitochondria damaged by irradiation may persist in the cytoplasm and retain partial respiratory activity, and simultaneous inhibition of autophagy and mitophagy in the presence of metabolically active, damaged mitochondria could further enhance radiosensitivity. In contrast, the minimal change in radiosensitivity following ATG2A knockdown may reflect functional redundancy with ATG2B. Alternatively, the increase in AR expression observed after ATG2A knockdown may have partially offset any pro-apoptotic effects of ATG2A loss. Taken together, these observations raise the possibility of a regulatory interaction between ATG2A and AR signaling.

In this study, we used enzalutamide and abiraterone acetate as AR inhibitors. Enzalutamide inhibits the nuclear translocation of AR, and abiraterone inhibits the activity of 5α-reductase, which converts testosterone to DHT. However, in our experiment, while AR activation by DHT was suppressed by both abiraterone and enzalutamide, abiraterone exhibited a clearer radiosensitizing effect, consistent with previous reports.

As a secondary aim, we explored the impact of metabolism on radiation response by regulating glucose availability. Several previous studies have reported that glucose concentrations and intermittent fasting may influence radiosensitivity, with findings indicating that the MAPK pathway, including p38, promotes DNA repair and helps maintain the integrity of normal tissues following radiation exposure (48,49). Our results showed that, while normal cells do not typically undergo autophagy in response to radiation, they may activate AMPK and p38 MAPK under LG conditions, potentially accelerating DNA repair. In contrast, AR-positive PCa cell lines did not exhibit any changes. Our findings suggest that glucose restriction or fasting during PCa radiotherapy might help protect radiosensitive pelvic tissues, such as the intestines, kidneys, and bladder, by enhancing the DNA repair capacity and stress response of normal cells without conferring the same protective effect on AR-positive tumor cells.

## CRediT authorship contribution statement

**K.K.:** Writing – original draft, Writing – reviewing & editing, Conceptualization, Methodology, Data curation, Investigation, Project administration, Funding acquisition. **Q.F.:** Methodology, Investigation, Data curation. **M.I.C.:** Methodology, Investigation, Data curation. **K.A.:** Writing – reviewing & editing. **J.M.A.:** Writing – original draft, Writing – reviewing & editing, Supervision. **L.W.:** Writing – reviewing & editing. **Z.H.:** Resources. **J.M.:** Methodology, Investigation, Data curation. **M.M.:** Methodology, Investigation. **E.S.** Methodology, Investigation. **Z.V.W.:** Resources, Methodology, Conceptualization, Writing – reviewing & editing, Supervision. **J.R.:** Writing – reviewing & editing, Supervision. **Y.L.:** Writing – reviewing & editing, Supervision, Project administration, Funding acquisition.

## Acknowledgements

We thank Drs. Brian Armstrong and Martha Salas at the City of Hope Light Microscopy Core Facility for their assistance with confocal microscopy; Drs. Zhuo Li and Ricardo Zerda at the City of Hope Electron Microscopy Core Facility for their help with transmission electron microscopy; Professor Jose Enrique Montero Casimiro at the City of Hope Arthur Riggs Diabetes & Metabolism Research Institute for fluorescent microscopy; Dr. Alan Ly, Scientific Writer for the Department of Radiation Oncology at the City of Hope Comprehensive Cancer Center for assistance with editing the manuscript; Dr. Sarah Shuck at the Department of Diabetes & Cancer Metabolism for providing the research materials; Dr. Jinhui Wang at the City of Hope Integrative Genomics Core Facility for conducting RNA-seq experiments; Dr. Terence Williams’ lab and Dr. Binghui Shen’s lab for providing experimental equipment; and all the staff members in the City of Hope Comprehensive Cancer Center for their kind support

## SUPPLEMENTARY INFORMATION

### Supplementary Figure legends

**Supplementary table 1.** Primary and secondary antibody list.

| Antigen | Clone | Supplier | Conjugation | Catalog no. | Specie | Experiment | Conc. |
| --- | --- | --- | --- | --- | --- | --- | --- |
| Primary antibodies |  |  |  |  |  |  |  |
| LC3B | polyclonal | Thermo Fisher | unconjugate | PA1-46286 | Rb | WB, IHC-F | 1:1000, |
| CK8 | EP1628Y | Abcam | unconjugate | 53280 | Rb | IHC | 1:200 |
| ATG9A | EPR2450( | Abcam | unconjugate | 108338 | Rb | IHC-F | 1:50 |
| AR | AR441 | Abcam | unconjugate | 9474 | Ms | IHC-F | 1:50 |
| CD326 | VU-1D9 | Bio-Rad | unconjugate | MCA1870G | Ms | ICC | 1:200 |
| Phospho-AMPKα | - | Cell Signaling | unconjugate | 2531 | Rb | WB | 1:1000 |
| GAPDH | 14C10 | Cell Signaling | unconjugate | 2118 | Rb | WB | 1:1000 |
| Isotype control for | DA1E | Cell Signaling | AF488 | 2975 | Rb | FCM | 1:100 |
| Cyclin D1 | 92G2 | Cell Signaling | unconjugate | 2978 | Rb | WB | 1:1000 |
| Isotype control for CC3 | DA1E | Cell Signaling | AF647 | 2985 | Rb | FCM | 1:100 |
| Phospho-Akt (Ser473) | D9E | Cell Signaling | unconjugate | 4060 | Rb | WB | 1:2000 |
| Phospho-p38 | D3F9 | Cell Signaling | unconjugate | 4511 | Rb | WB | 1:1000 |
| CK8/18 | C51 | Cell Signaling | unconjugate | 4546 | Ms | IHC | 1:50 |
| βActin | 13E5 | Cell Signaling | unconjugate | 4970 | Rb | WB | 1:1000 |
| AR | D6F11 | Cell Signaling | unconjugate | 5153 | Rb | IHC | 1:800 |
| PSA | D6B1 | Cell Signaling | unconjugate | 5365 | Rb | IHC | 1:400 |
| Phospho-Rb | D20B12 | Cell Signaling | unconjugate | 8516 | Rb | WB | 1:1000 |
| Rad51 | D4B10 | Cell Signaling | unconjugate | 8875 | Rb | WB | 1:1000 |
| Cleaved Caspase-3 | D3E9 | Cell Signaling | AF647 | 9602 | Rb | FCM | 1:100 |
| Cleaved Caspase-3 | D3E9 | Cell Signaling | unconjugate | 9661 | Rb | IHC-F, ICC | 1:200, 1:800 |
| Phospho-H2A.X | 20E3 | Cell Signaling | unconjugate | 9718 | Rb | IHC-F, ICC | 1:200 |
| Phospho-H2A.X | 20E3 | Cell Signaling | AF488 | 9719 | Rb | FCM | 1:100 |
| AR-V7 | E3O8L | Cell Signaling | unconjugate | 19672 | Rb | WB | 1:1000 |
| DNA-PKcs | E6U3A | Cell Signaling | unconjugate | 38168 | Rb | WB | 1:1000 |
| Phospho-DNA-PKcs | E9J4G | Cell Signaling | unconjugate | 68716 | Rb | WB | 1:1000 |
| Secondary antibodies |  |  |  |  |  |  |  |
| IgG |  | Cell Signaling | unconjugate | 7076 | Ms | WB | 1:3000-5000 |
| IgG |  | Cell Signaling | unconjugate | 7074 | Rb | WB | 1:3000-5000 |
| IgG |  | abcam | AF555 | 150114 | Ms | IHC-F/ICC | 1:200 |
| IgG |  | ThermoFisher | AF488 | A11008 | Rb | IHC-F/ICC | 1:500 |

Western blot: WB, Flow Cytometry : FCM, Fluorescent immunohistochemistry: IHC-F, immunocytochemistry: ICC, Ms: Mouse, Rb: Rabbit,

**Supplementary table 2.** Products used in siRNA knockdown experiment.

| siRNA targeting gene | Target exon | Product company | Cat# |
| --- | --- | --- | --- |
| *AR* | Exon 2 | ThermoFisher Scientific | 4390824, s1538 |
| *AR* | Exon 7 |  | 4390824, s1539 |
| *ATG2A* | Exon 32 |  | 4392420, s23097 |
| *BECN1* | Exon 5-7 |  | 4392420, s16537 |
| *MAP1LC3B* | Exon 2 and 3 |  | 4392420, s37749 |

**Supplementary table 3.**
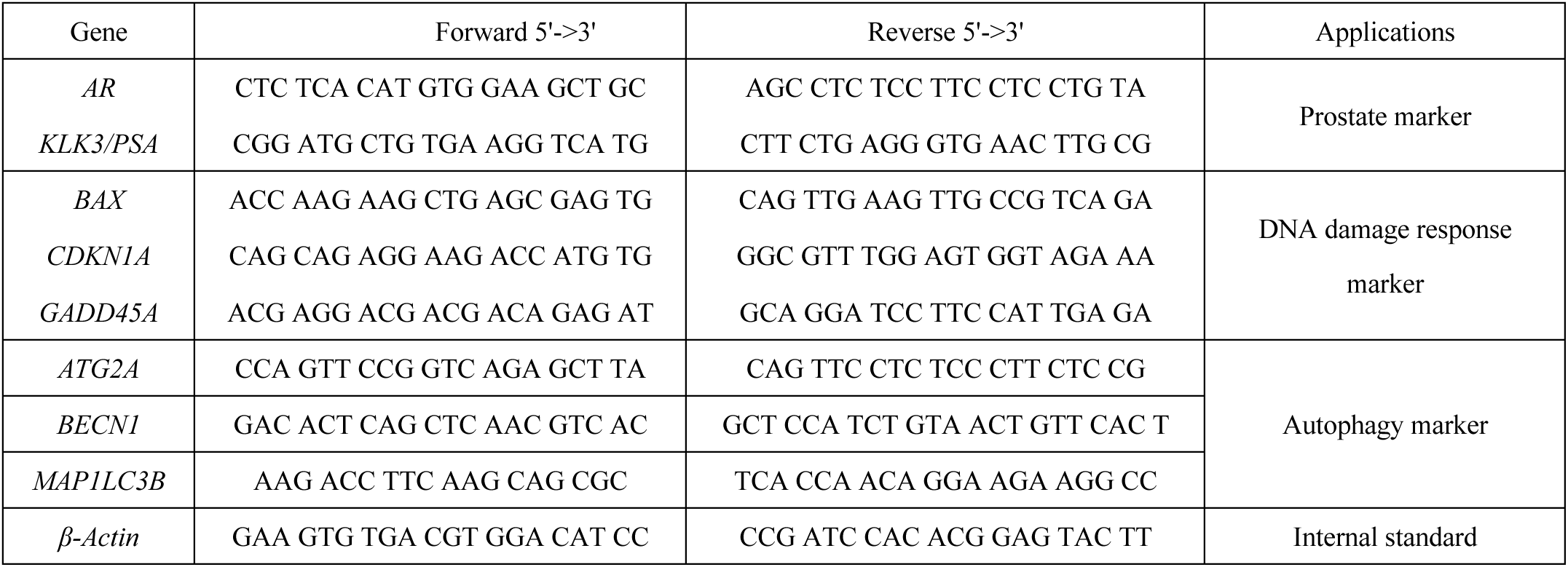
Sequence of primers used in RT-qPCR.

**Supplementary Figure 1.**
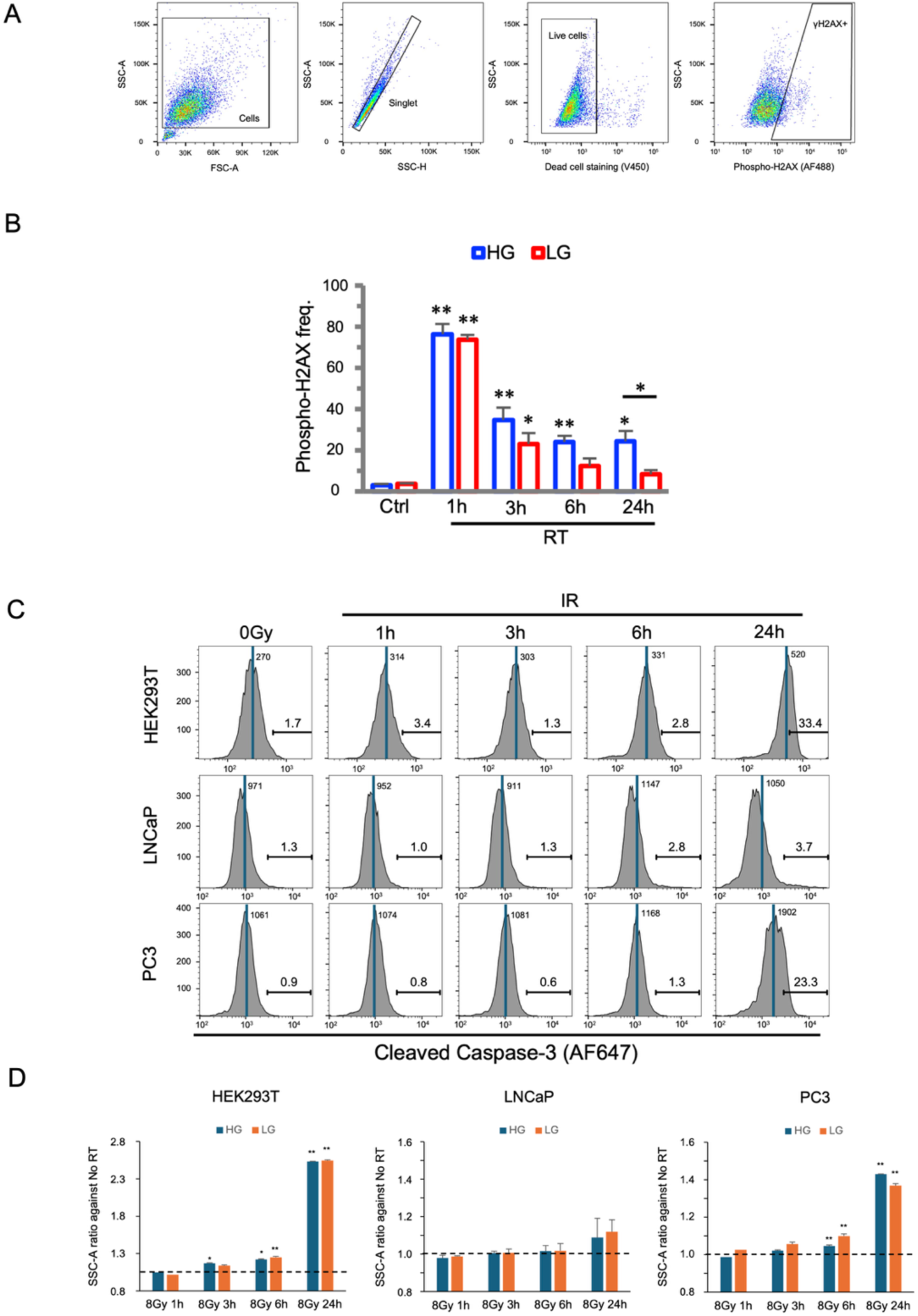
**A,** The plots of flow cytometry from selection of cells to detection of γH2AX signals. **B,** Comparison of γH2AX frequencies generated in HEK293T between high glucose (HG) and low glucose (LG) during 24 hours after irradiation. **C,** Representative profiles for cleaved caspase-3 (CC3) frequencies in HEK293T, LNCaP, and PC3. The numbers listed in each panel mean the median value. **D,** Side scatter (SSC) ratio calculated from SSC-A values in irradiated and unirradiated groups. *, P<0.05, **, P<0.01 vs. 0Gy.

**Supplementary Figure 2.**
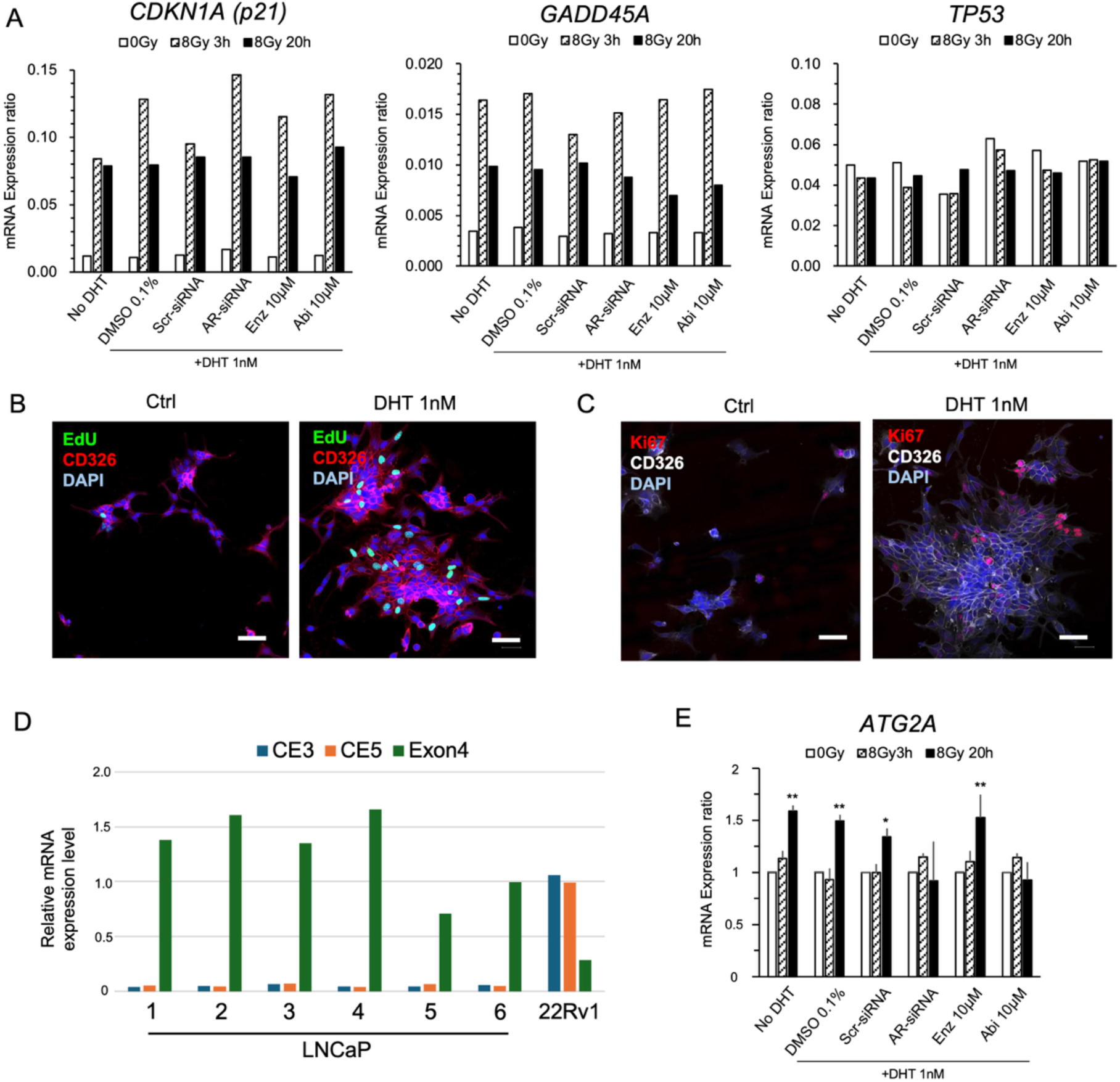
**A,** mRNA expression levels of *CDKN1A*, *GADD45A*, and *TP53* genes in LNCaP cells. Enz and Abi mean enzalutamide and abiraterone acetate. Data represent the mean values of two RT-qPCR assays. **B** and **C,** ICC images of LNCaP cells stained with (**B**) fluorescent azide targeting EdU incorporated in DNA strands and (**C**) anti-Ki67 antibody. **D,** mRNA expression levels of Exon 4 and cryptic exons (*CE3* and *CE5*) in AR gene. No. 1-6 in graph show the lot number of LNCaP cells. **E,** Relative mRNA expression levels of *ATG2A* gene in LNCaP cells. Data represent the means and SEs of three independent experiments. *, P<0.05, **, P<0.01.

**Supplementary Figure 3.**
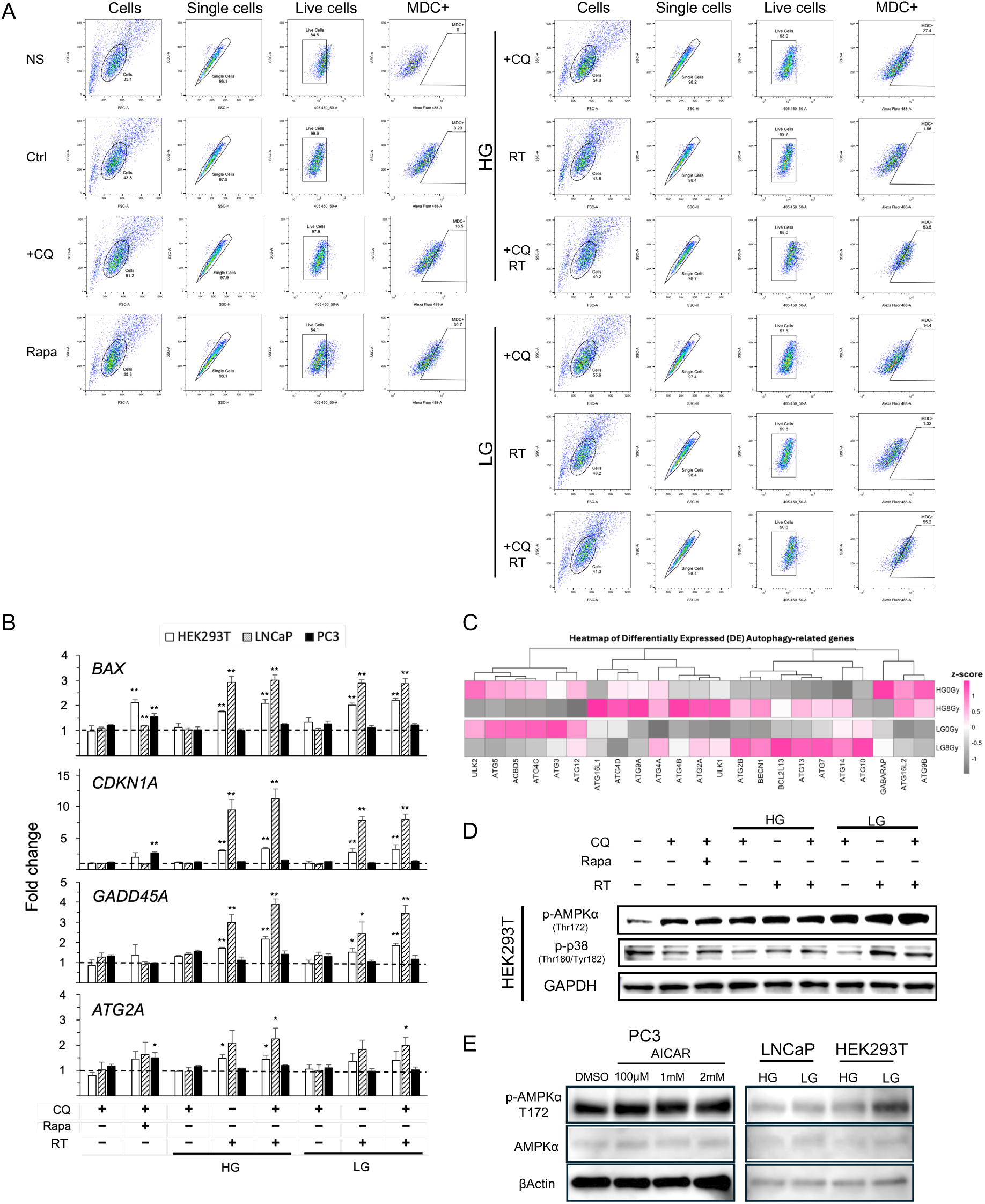
**A,** MDC profiles got with FCM. **B,** mRNA expression levels of p53-downstream genes in HEK293T, LNCaP, and PC3. The cells were treated in the same manner as MDC assay. The data show the fold change of each mRNA expression against negative control (CQ**−**, Rapa**−**, IR**−**). **C,** Heatmap showing differentially expressed autophagy-related gene expression in unirradiated (0 Gy) and irradiated (8 Gy). LNCaP cells were cultured in RPMI1640 containing 25 mM glucose (HG) or 5 mM glucose (LG). **D,** Phosphorylated AMPKα (Thr172; p-AMPKα) and dual phosphorylated p38 (Thr180/Tyr182; p-p38) were measured with western blot analysis, which are representative markers of mitogen-activated protein kinases (MAPK). **E,** p-AMPKα and AMPKα measured in PC3, LNCaP and HEK293T. GAPDH or βActin was used as an internal control.

**Supplementary Figure 4.**
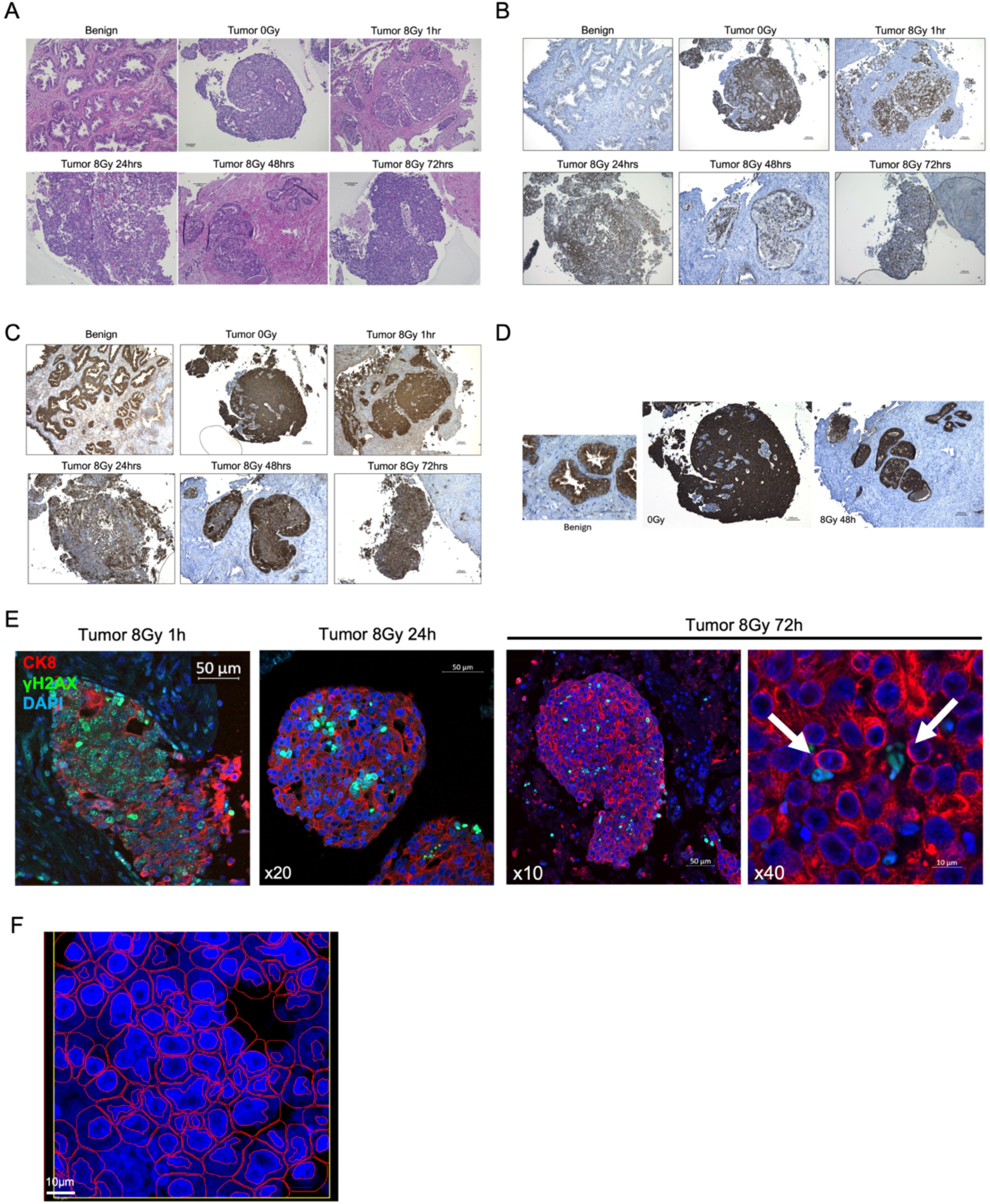
IHC images of patient-derived explants. (**A**) H&E staining images, (**B**) AR, (**C**) PSA, and (**D**) CK8 stained by DAB. (**E**) IHC-F of CK8 and γH2AX after irradiation. (**F**) Representative image showing QuPath-based cell segmentation in PCa tissue. Nuclei were stained with DAPI (blue), and cell segmentation was performed using the DAPI channel as a reference in QuPath. Detected cell boundaries are outlined in red.

## References

1. Siegel RL, Miller KD, Wagle NS, Jemal A. Cancer statistics, 2023. CA Cancer J Clin 2023;73:17–48

2. Germain L, Lafront C, Paquette V, Neveu B, Paquette JS, Pouliot F, et al. Preclinical models of prostate cancer - modelling androgen dependency and castration resistance in vitro, ex vivo and in vivo. Nat Rev Urol 2023;20:480–93

3. Audet-Walsh E, Yee T, McGuirk S, Vernier M, Ouellet C, St-Pierre J, et al. Androgen-Dependent Repression of ERRgamma Reprograms Metabolism in Prostate Cancer. Cancer Res 2017;77:378–89

4. Loblaw DA, Virgo KS, Nam R, Somerfield MR, Ben-Josef E, Mendelson DS, et al. Initial hormonal management of androgen-sensitive metastatic, recurrent, or progressive prostate cancer: 2006 update of an American Society of Clinical Oncology practice guideline. J Clin Oncol 2007;25:1596–605

5. Sturge J, Caley MP, Waxman J. Bone metastasis in prostate cancer: emerging therapeutic strategies. Nat Rev Clin Oncol 2011;8:357–68

6. Zhang W, van Gent DC, Incrocci L, van Weerden WM, Nonnekens J. Role of the DNA damage response in prostate cancer formation, progression and treatment. Prostate Cancer Prostatic Dis 2020;23:24–37

7. Tarish FL, Schultz N, Tanoglidi A, Hamberg H, Letocha H, Karaszi K, et al. Castration radiosensitizes prostate cancer tissue by impairing DNA double-strand break repair. Sci Transl Med 2015;7:312re11

8. Goodwin JF, Schiewer MJ, Dean JL, Schrecengost RS, de Leeuw R, Han S, et al. A hormone-DNA repair circuit governs the response to genotoxic insult. Cancer Discov 2013;3:1254–71

9. Polkinghorn WR, Parker JS, Lee MX, Kass EM, Spratt DE, Iaquinta PJ, et al. Androgen receptor signaling regulates DNA repair in prostate cancers. Cancer Discov 2013;3:1245–53

10. Li L, Karanika S, Yang G, Wang J, Park S, Broom BM, et al. Androgen receptor inhibitor-induced “BRCAness” and PARP inhibition are synthetically lethal for castration-resistant prostate cancer. Sci Signal 2017;10

11. Jang M, Park R, Kim H, Namkoong S, Jo D, Huh YH, et al. AMPK contributes to autophagosome maturation and lysosomal fusion. Sci Rep 2018;8:12637

12. Kim J, Kundu M, Viollet B, Guan KL. AMPK and mTOR regulate autophagy through direct phosphorylation of Ulk1. Nat Cell Biol 2011;13:132–41

13. Wang W, Liu J, Wu Q. MiR-205 suppresses autophagy and enhances radiosensitivity of prostate cancer cells by targeting TP53INP1. European Review for Medical & Pharmacological Sciences 2016;20

14. Blessing AM, Rajapakshe K, Reddy Bollu L, Shi Y, White MA, Pham AH, et al. Transcriptional regulation of core autophagy and lysosomal genes by the androgen receptor promotes prostate cancer progression. Autophagy 2017;13:506–21

15. Guo JY, Teng X, Laddha SV, Ma S, Van Nostrand SC, Yang Y, et al. Autophagy provides metabolic substrates to maintain energy charge and nucleotide pools in Ras-driven lung cancer cells. Genes Dev 2016;30:1704–17

16. Roy A, Bera S, Saso L, Dwarakanath BS. Role of autophagy in tumor response to radiation: Implications for improving radiotherapy. Frontiers in oncology. Volume 122022. p 957373.

17. White MA, Tsouko E, Lin C, Rajapakshe K, Spencer JM, Wilkenfeld SR, et al. GLUT12 promotes prostate cancer cell growth and is regulated by androgens and CaMKK2 signaling. Endocr Relat Cancer 2018;25:453–69

18. Lin C, Blessing AM, Pulliam TL, Shi Y, Wilkenfeld SR, Han JJ, et al. Inhibition of CAMKK2 impairs autophagy and castration-resistant prostate cancer via suppression of AMPK-ULK1 signaling. Oncogene 2021;40:1690–705

19. Frigo DE, Howe MK, Wittmann BM, Brunner AM, Cushman I, Wang Q, et al. CaM kinase kinase beta-mediated activation of the growth regulatory kinase AMPK is required for androgen-dependent migration of prostate cancer cells. Cancer Res 2011;71:528–37

20. Mascorro JA, Bozzola JJ. Processing biological tissues for ultrastructural study. Methods Mol Biol 2007;369:19–34

21. Wlodkowic D, Telford W, Skommer J, Darzynkiewicz Z. Apoptosis and beyond: cytometry in studies of programmed cell death. Methods Cell Biol 2011;103:55–98

22. Ding D, Zhang Y, Wang J, Zhang X, Gao Y, Yin L, et al. Induction and inhibition of the pan-nuclear gamma-H2AX response in resting human peripheral blood lymphocytes after X-ray irradiation. Cell Death Discov 2016;2:16011

23. Davis ID, Stockler MR, Sweeney CJ. Enzalutamide in Metastatic Prostate Cancer. Reply. N Engl J Med 2019;381:1494–5

24. Wright TC, Dunne VL, Alshehri AHD, Redmond KM, Cole AJ, Prise KM. Abiraterone In Vitro Is Superior to Enzalutamide in Response to Ionizing Radiation. Front Oncol 2021;11:700543

25. Balk SP, Knudsen KE. AR, the cell cycle, and prostate cancer. Nucl Recept Signal 2008;6:e001

26. Helin K, Harlow E, Fattaey A. Inhibition of E2F-1 transactivation by direct binding of the retinoblastoma protein. Mol Cell Biol 1993;13:6501–8

27. Sherr CJ, Roberts JM. CDK inhibitors: positive and negative regulators of G1-phase progression. Genes Dev 1999;13:1501–12

28. Cobrinik D. Pocket proteins and cell cycle control. Oncogene 2005;24:2796–809

29. Mizushima N, Yoshimori T. How to interpret LC3 immunoblotting. Autophagy 2007;3:542–5

30. Biederbick A, Kern HF, Elsasser HP. Monodansylcadaverine (MDC) is a specific in vivo marker for autophagic vacuoles. Eur J Cell Biol 1995;66:3–14

31. Levine B, Kroemer G. Biological Functions of Autophagy Genes: A Disease Perspective. Cell 2019;176:11–42

32. Li X, He S, Ma B. Autophagy and autophagy-related proteins in cancer. Mol Cancer 2020;19:12

33. Klionsky DJ, Abdel-Aziz AK, Abdelfatah S, Abdellatif M, Abdoli A, Abel S, et al. Guidelines for the use and interpretation of assays for monitoring autophagy (4th edition)(1). Autophagy 2021;17:1–382

34. Karabiyik C, Vicinanza M, Son SM, Rubinsztein DC. Glucose starvation induces autophagy via ULK1-mediated activation of PIKfyve in an AMPK-dependent manner. Dev Cell 2021;56:1961–75 e5

35. Choi SY, Kim MJ, Kang CM, Bae S, Cho CK, Soh JW, et al. Activation of Bak and Bax through c-abl-protein kinase Cdelta-p38 MAPK signaling in response to ionizing radiation in human non-small cell lung cancer cells. J Biol Chem 2006;281:7049–59

36. Zarubin T, Han J. Activation and signaling of the p38 MAP kinase pathway. Cell Res 2005;15:11–8

37. Ponguta LA, Gregory CW, French FS, Wilson EM. Site-specific androgen receptor serine phosphorylation linked to epidermal growth factor-dependent growth of castration-recurrent prostate cancer. J Biol Chem 2008;283:20989–1001

38. Chymkowitch P, Le May N, Charneau P, Compe E, Egly JM. The phosphorylation of the androgen receptor by TFIIH directs the ubiquitin/proteasome process. Embo j 2011;30:468–79

39. Rahman R, Selth LA. Cyclin-dependent kinases as mediators of aberrant transcription in prostate cancer. Transl Oncol 2025;55:102378

40. Samaranayake GJ, Troccoli CI, Huynh M, Lyles RDZ, Kage K, Win A, et al. Thioredoxin-1 protects against androgen receptor-induced redox vulnerability in castration-resistant prostate cancer. Nat Commun 2017;8:1204

41. Spratt DE, Evans MJ, Davis BJ, Doran MG, Lee MX, Shah N, et al. Androgen Receptor Upregulation Mediates Radioresistance after Ionizing Radiation. Cancer Res 2015;75:4688–96

42. Kang R, Zeh HJ, Lotze MT, Tang D. The Beclin 1 network regulates autophagy and apoptosis. Cell Death Differ 2011;18:571–80

43. Jia J, Zhang HB, Shi Q, Yang C, Ma JB, Jin B, et al. KLF5 downregulation desensitizes castration-resistant prostate cancer cells to docetaxel by increasing BECN1 expression and inducing cell autophagy. Theranostics 2019;9:5464–77

44. Zhang M, Sun Y, Meng J, Zhang L, Liang C, Chang C. Targeting AR-Beclin 1 complex-modulated growth factor signaling increases the antiandrogen Enzalutamide sensitivity to better suppress the castration-resistant prostate cancer growth. Cancer Lett 2019;442:483–90

45. Kon N, Ou Y, Wang SJ, Li H, Rustgi AK, Gu W. mTOR inhibition acts as an unexpected checkpoint in p53-mediated tumor suppression. Genes Dev 2021;35:59–64

46. Fernandez AF, Sebti S, Wei Y, Zou Z, Shi M, McMillan KL, et al. Disruption of the beclin 1-BCL2 autophagy regulatory complex promotes longevity in mice. Nature 2018;558:136–40

47. Lazarou M, Sliter DA, Kane LA, Sarraf SA, Wang C, Burman JL, et al. The ubiquitin kinase PINK1 recruits autophagy receptors to induce mitophagy. Nature 2015;524:309–14

48. Bengal E, Aviram S, Hayek T. p38 MAPK in Glucose Metabolism of Skeletal Muscle: Beneficial or Harmful? Int J Mol Sci 2020;21

49. de la Cruz Bonilla M, Stemler KM, Jeter-Jones S, Fujimoto TN, Molkentine J, Asencio Torres GM, et al. Fasting Reduces Intestinal Radiotoxicity, Enabling Dose-Escalated Radiation Therapy for Pancreatic Cancer. Int J Radiat Oncol Biol Phys 2019;105:537–47

